# Durable suppression of viremia by lipid nanoparticle-formulated RNA encoding for a highly potent HIV-1 neutralizing antibody

**DOI:** 10.64898/2026.09.17.752120

**Authors:** Sophie Sayettat, Felix Tolksdorf, Robin Johannson, Johannes Nelke, Alexandra Malz, Leyla Fischer, Imke Gerhard, Nathalie Ullrich, Ursula Ellinghaus, Christiane R. Stadler, Jan P. Bogen, Candice Morin, Jacqueline Knüfer, Ricarda Stumpf, Philipp Schommers, Uğur Şahin, Michael S. Seaman, Sven Kratochvil, Florian Klein, Valentin Le Douce, Henning Gruell

## Abstract

Recombinant broadly neutralizing antibodies (bNAbs) are promising tools to treat and prevent HIV-1 infection but are associated with manufacturing challenges. To overcome this limitation, we encoded the potent bNAb 1-18-LS on RNA (1-18-LS RibobNAb) for delivery via lipid nanoparticles (LNPs). Indicating high *in vivo* antibody expression, a 30 µg RNA-LNP intravenous injection resulted in an average peak 1-18-LS RibobNAb serum concentration of 1,061 µg/mL in human neonatal Fc receptor-transgenic mice. Notably, weekly 30 µg RNA-LNP doses maintained higher trough bNAb levels than 500 µg protein injections. Highlighting potent antiviral activity, RNA-LNP-mediated 1-18-LS RibobNAb monotherapy of viremic HIV-1_YU2_-infected humanized mice resulted in durable HIV-1 suppression without emerging viral escape. Importantly, 1-18-LS RNA-LNP treatment also fully controlled infection after interruption of antiretroviral therapy in humanized mice infected with different patient-derived polyclonal HIV-1 isolates. Our findings provide a proof-of-principle for effective RNA-mediated bNAb immunotherapy of HIV-1 infection.

## INTRODUCTION

Potent neutralizing antiviral antibodies are effective in the treatment and prevention of infectious diseases including Ebola virus disease, Covid-19, and severe RSV infection.^1,2^ In the clinic, monoclonal antibodies are typically administered by infusion or injection of recombinant protein produced in mammalian cell lines. Complex manufacturing processes and requirements for extensive purification can result in challenges for large-scale production, high costs, and limited availability of antibody-based drugs.^3,4^ By facilitating endogenous antibody expression, delivery of antibody-encoding RNA represents a potential alternative to the use of recombinant proteins.^5–9^ Proof-of-concept studies in animal models demonstrated that administration of antibody-encoding RNA can protect against infection with viruses including SARS-CoV-2, RSV, chikungunya virus (CHIKV), and HIV-1.^10–14^ However, whether RNA-based antibody therapeutics are effective in treating active viral infection has not been established.

While pathogen resistance poses a potential problem for all antibody-based strategies, targeting the HIV-1 envelope protein (Env) is particularly challenging due to its high diversity and mutability.^15^ Nevertheless, broadly neutralizing antibodies (bNAbs) with activity against a large fraction of HIV-1 strains have been isolated from people living with HIV.^16^ Amongst these bNAbs, antibodies targeting the HIV-1 Env CD4 binding site (CD4bs), which has a critical function in the viral replication cycle, represent some of the most potent and broad candidates identified to date.^17–24^ Clinical trials demonstrated that CD4bs bNAbs can reduce plasma viral loads, delay viral rebound after ART interruption, or prevent infection with highly sensitive viral strains.^25–32^ However, although tolerance for variation in the CD4bs may be limited and antibody escape can be associated with reduced viral fitness,^33–36^ clinical effects of CD4bs bNAbs were constrained by pre-existing or emerging antibody resistance.^25,26,32^ Thus, bNAbs with higher activity may be required for optimal antibody-based strategies against HIV-1. Compared with previously isolated bNAbs, the potent CD4bs antibody 1-18 is notable for its limited HIV-1 escape identified *in vitro* and *in vivo*.^17^ The half-life-optimized 1-18-LS variant is under clinical investigation as a recombinant protein (also referred to as BNT351).^37^

Here we combine RNA-LNP delivery technology with bNAb 1-18-LS to develop an effective antibody-based strategy against HIV-1 that is independent of recombinantly produced antibody. Using a combination of *in vitro* and *in vivo* experiments, we demonstrate RNA-mediated bNAb expression with a favorable pharmacokinetic profile and durable suppression of viremia extending to clinical HIV-1 isolates. Our results provide a proof-of-principle for effective RNA-based immunotherapy of HIV-1 infection.

## RESULTS

### Potent HIV-1 neutralization by 1-18-LS RibobNAb

To achieve effective antibody expression, we generated 1-methylpseudouridine (m1Ψ)-modified RNA encoding heavy and light chains of 1-18 (referred to as RibobNAb when expressed from RNA constructs) based on the RiboMab platform (**Figure 1A**).^7^ To optimize antibody half-life, we generated the 1-18-LS variant incorporating Fc domain substitutions M428L and N434S that increase affinity to the human neonatal Fc receptor (hFcRn) at pH 6.0.^38^ For effective *in vivo* delivery of RNA, heavy and light chain RNA molecules were co-formulated at a 1.5:1.0 µg ratio into lipid nanoparticles (LNPs) (**Figure 1A**).

**Figure 1.**
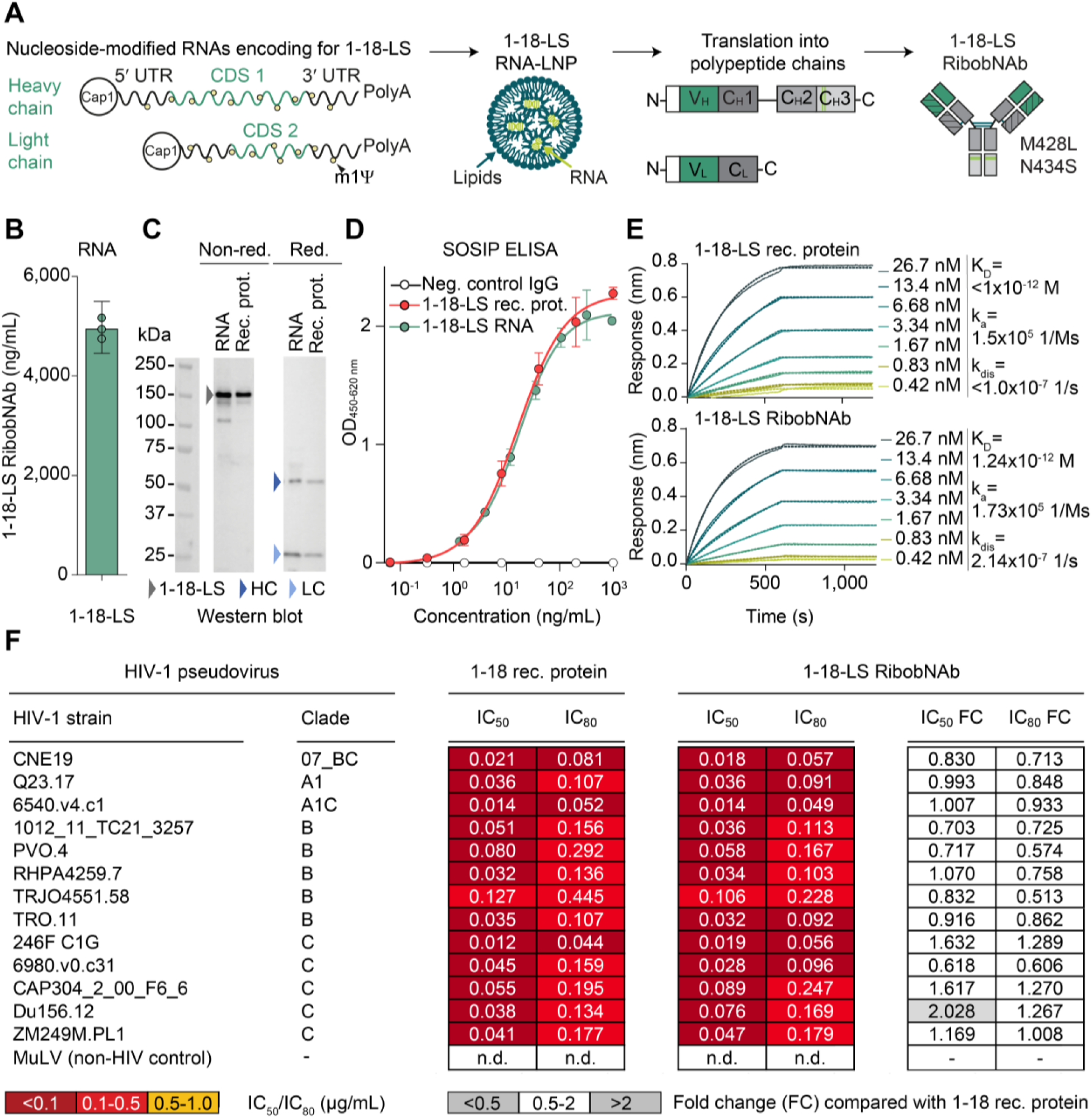
1-18-LS RibobNAb potently neutralizes HIV-1. **(A)** Nucleoside-modified RNA sequences encoding for 1-18-LS heavy or light chain are co-formulated into lipid nanoparticles (LNPs), resulting in expression of 1-18-LS RibobNAb upon cellular uptake and translation. m1Ψ, 1-methylpseudouridine; UTR, untranslated region; CDS, coding DNA sequence. **(B)** 1-18-LS RibobNAb levels in HEK293T/17 cell culture supernatant after heavy and light chain RNA co-transfection. Circles indicate different transfections; bar and error bar show geometric mean and 95% confidence interval, respectively. (**C**) Western blot analysis of 1-18-LS RibobNAb antibody chain expression in RNA-transfected HEK293T/17 cell culture supernatants compared with recombinant 1-18-LS protein (rec. prot.). Left, center, and right panels show molecular weight ladder, non-reducing (non-red.), and reducing conditions, respectively. Arrows indicate 1-18-LS antibody, heavy chain (HC), and light chain (LC). (**D**) HIV-1 Env BG505 DS-SOSIP.664 ELISA of 1-18-LS recombinant protein and 1-18-LS RibobNAb in RNA-transfected HEK293T/17 cell culture supernatants. Symbols and error bars indicate arithmetic means and standard deviations, respectively (representative experiment with technical duplicates for RibobNAb; recombinant protein tested in four replicates). (**E**) HIV-1 Env BG505 DS-SOSIP.664 biolayer interferometry measurement of recombinant 1-18-LS protein and 1-18-LS RibobNAb in RNA-transfected HEK293T/17 cell culture supernatant. K_D_, dissociation constant; k_a_, association rate constant; k_dis_, dissociation rate constant. (**F**) HIV-1 pseudovirus neutralization of 1-18 recombinant protein and 1-18-LS RibobNAb in RNA-transfected HEK293T/17 cell culture supernatants. For supernatants, values represent mean IC_50_s and IC_80_s of triplicates. n.d., not detected at minimum concentrations of 0.30 µg/mL (1-18 protein) or 0.28 µg/mL (1-18-LS RibobNAb). FC, fold change.

Expression of 1-18-LS and 1-18 from RNA was confirmed in supernatants of transfected 293T cells (geometric mean concentrations of 4,948 and 4,192 ng/mL for 1-18-LS and 1-18, respectively) (**Figures 1B** and **1C**; **Figures S1A** and **S1B**). Enzyme-linked immunosorbent assay (ELISA) and biolayer interferometry (BLI) demonstrated potent 1-18-LS RibobNAb binding and high affinity to the near-native prefusion closed state-stabilized HIV-1 BG505 DS-SOSIP.664 Env trimer, respectively (half-maximal effective concentration [EC_50_] of 0.014 µg/mL and K_d_ of 1.24x10^-12^ M) (**Figures 1D** and **1E**; **Figure S1C**).^39^ To confirm that the RibobNAb format retains the high neutralizing activity of 1-18, we performed TZM-bl cell-based HIV-1 neutralization assays using RNA-transfected cell culture supernatants against panels of up to 16 pseudovirus strains representing diverse HIV-1 clades. Pseudovirus neutralization of 1-18-LS RibobNAb and 1-18 RibobNAb strongly correlated (Pearson’s *r*=0.99, p<0.001; based on IC_50_s for the ten pseudoviruses tested against both constructs) (**Figure 1F**; **Figures S1D** and **S1E**). Importantly, both RibobNAb constructs showed potent neutralizing activity that was near-identical to that of recombinant 1-18 protein (geometric mean fifty-percent inhibitory concentrations [GeoMean IC_50_s] of 0.05 µg/ml against the tested viral strains; average 1.13- and 1.20-fold IC_50_ changes relative to 1-18 recombinant protein, respectively) (**Figure 1F**; **Figures S1D** and **S1E**).

Thus, nucleoside-modified heavy and light chain RNA molecules enable effective expression of highly potent 1-18-LS RibobNAb. Importantly, analysis of 293T cell supernatants confirmed expression of functional 1-18-LS RibobNAb from lipid nanoparticle-formulated RNA (**Figures S1C**, **S1F** and **S1G**), supporting the use of RNA-LNP constructs for *in vivo* delivery.

### RNA-LNP administration leads to high 1-18-LS RibobNAb levels *in vivo*

To investigate RNA-LNP-mediated 1-18-LS RibobNAb expression in an *in vivo* model relevant for predicting human pharmacokinetics, we determined systemic antibody levels in hFcRn-transgenic mice (**Figure S2A**).^40,41^ Serum analysis after injection of LNP-formulated RNA indicated correct antibody assembly without signs of aggregate formation (**Figure S2B**). Following an intravenous administration of 10 µg or 30 µg RNA-LNP, peak average 1-18-LS RibobNAb serum levels of 322 and 1,061 µg/mL were observed within two to three days, respectively (**Figure 2A**). In contrast to the rapid decline of circulating antibody to undetectable levels after administration of 100 µg 1-18-LS protein, 1-18-LS RibobNAb remained detectable for ≥4 weeks and average levels >10 µg/mL were maintained for ≥32 days after a 30 µg RNA-LNP injection (**Figure 2A**). 1-18-LS RibobNAb expression was confirmed in hFcRn-negative NRG mice commonly used in mouse models of HIV-1 infection (**Figure 2B**).^42^ While antibody levels in these mice were lower than in hFcRn-transgenic mice, 1-18-LS RibobNAb serum concentrations >10 µg/mL were maintained for ≥14 days after a single 30 µg RNA-LNP injection and were overall higher when compared to a 200 µg 1-18-LS protein dose (**Figure 2B**). Potent and HIV-1-specific neutralizing serum activity correlating with antibody levels confirmed RNA-LNP-mediated expression of functional 1-18-LS RibobNAb (peak geometric mean fifty-percent inhibitory serum dilution [GeoMean ID_50_] of 11,102 against HIV-1 YU2 pseudovirus) (**Figure 2C**; **Figures S2C** and **S2D**).

**Figure 2.**
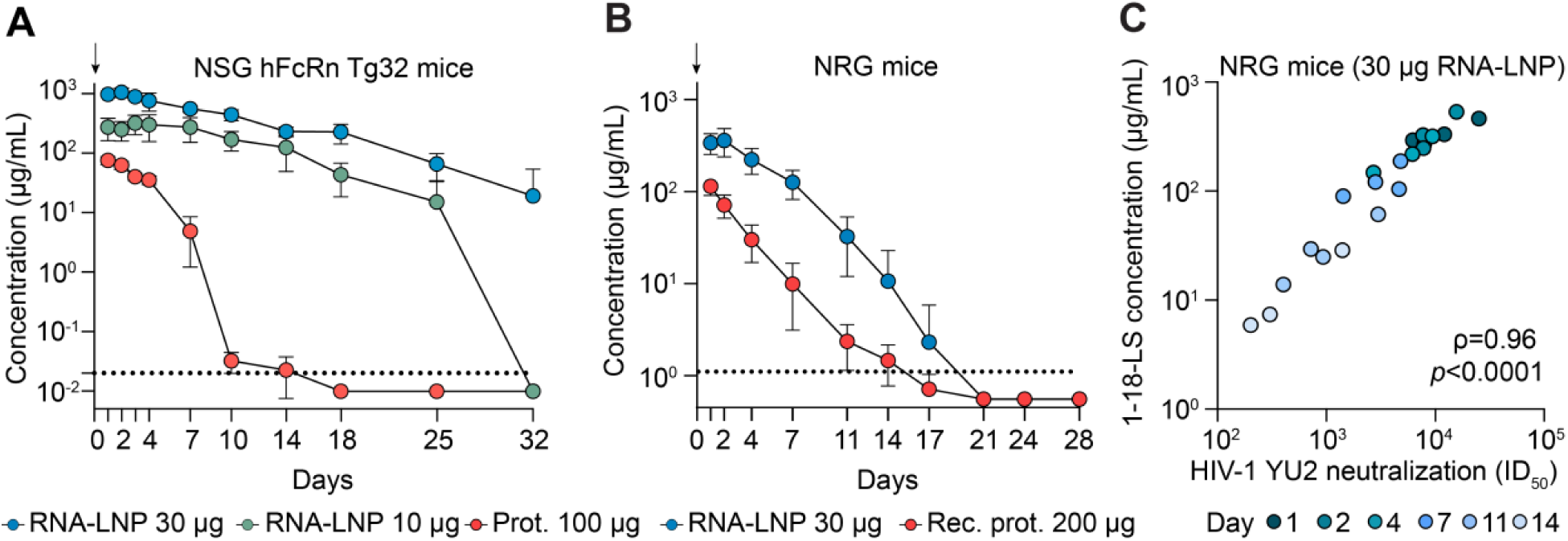
RNA-LNP leads to high *in vivo* 1-18-LS RibobNAb expression and potent HIV-1 neutralization. (**A**) Serum 1-18-LS levels in NSG hFcRn Tg32 mice after intravenous injection of RNA-LNP or recombinant protein. Symbols and error bars indicate arithmetic means and standard deviation *(n*=4). Dashed line indicates lower limit of quantification (0.02 µg/mL) and arrow represents injection. (**B**) Serum 1-18-LS levels in NRG mice after intravenous injection of RNA-LNP or recombinant protein. Symbols and error bars indicate arithmetic means and standard deviation (*n*=4). Dashed line indicates lower limit of quantification (1.11 µg/mL) and arrow represents injection. (**C**) Correlation of 1-18-LS RibobNAb serum concentrations and plasma HIV-1 YU2 pseudovirus neutralization ID_50_s longitudinally determined after an intravenous injection of 30 µg 1-18-LS RibobNAb RNA-LNP (*n*=4) in NRG mice. *ρ* indicates Spearman’s correlation coefficient.

As maintaining therapeutic antibody levels requires repeated dosing, we next investigated whether high *in vivo* RibobNAb expression persists in a setting of weekly i.v. 30 µg RNA-LNP injections. To this end, we longitudinally determined serum antibody levels and HIV-1 neutralizing activity in RNA-LNP-treated humanized NRG mice infected with HIV-1_YU2_. Mean trough 1-18-LS RibobNAb concentrations during weekly 30 µg RNA-LNP injections ranged from 45 to 82 µg/mL (range of 13 to 114 µg/mL in individual mice, respectively) (**Figure 3A**). Notably, throughout the seven-week treatment period, trough antibody concentrations were higher after administration of 30 µg 1-18-LS RNA-LNP when compared with weekly injections of 500 µg recombinant 1-18-LS protein (*p*=0.0003) (**Figure 3A**). Whereas HIV-1_YU2_-infected NRG mice did not show autologous activity against YU2 pseudovirus and RNA-LNP therapy did not induce unspecific neutralization of murine leukemia virus (MuLV) pseudovirus controls (**Figures S2E** and **S2F**), 1-18-LS RNA-LNP treatment resulted in potent HIV-1 serum neutralization (**Figure 3B**). Geometric mean ID_50_s in samples collected seven days after injection were numerically higher in mice receiving 30 µg 1-18-LS RibobNAb RNA-LNP than in mice receiving 500 µg 1-18-LS protein (GeoMean ID_50_s of 6,289 vs. 3,281 and 1,923 vs. 1,170 against HIV-1 YU2 and X2278 pseudoviruses, respectively) (**Figure 3B**). Although the differences in neutralization between RNA-LNP- and protein-treated mice were not statistically significant (*p*=0.18 and *p*=0.28), they were consistent with the higher 1-18-LS RibobNAb serum levels determined by anti-idiotype ELISA that strongly correlated with neutralizing activity (*ρ*=0.8 and *ρ*=0.9 for RNA-LNP and protein groups, respectively) (**Figures 3A-3C**).

**Figure 3.**
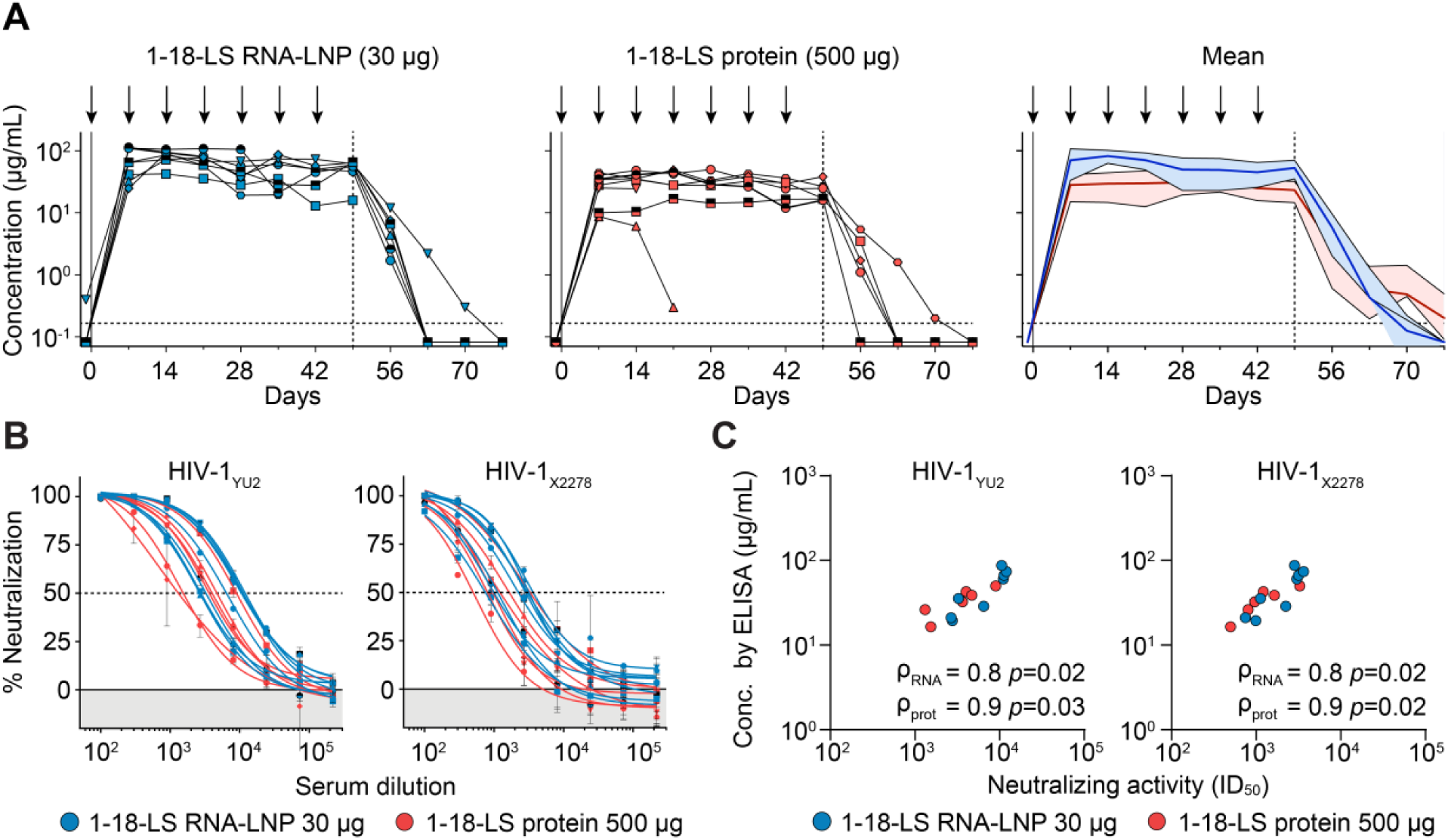
Weekly 1-18-LS RNA-LNP dosing sustains high RibobNAb *in vivo* expression. (**A**) Serum 1-18-LS trough levels in HIV-1_YU2_-infected CD34-humanized NRG mice. Right panel shows arithmetic means for both groups with standard deviation indicated by shaded areas (*n*=8). Arrows represent intravenous injections of RNA-LNP or recombinant protein. Vertical and horizontal dashed lines indicate end of treatment period and lower limit of quantification (0.165 µg/mL), respectively. (**B**) Neutralizing activity in HIV-1_YU2_-infected CD34-humanized NRG mouse serum collected seven days after RNA-LNP or recombinant protein injection against HIV-1 YU2 (left) and X2278 (right) pseudoviruses (collected on day 35 except for two protein group samples collected on days 28 or 42). Symbols and error bars indicate arithmetic means and standard deviation from two technical replicates, respectively (protein group, *n*=6; RNA-LNP group, *n*=8). Dashed line shows 50% neutralization. (**C**) Correlation of 1-18-LS levels and HIV-1 pseudovirus neutralizing activity (left, YU2 strain; right, X2278 strain) in HIV-1_YU2_-infected CD34-humanized NRG mouse serum collected seven days after RNA-LNP or recombinant protein injection (collected on day 35 except for two protein group samples collected on days 28 or 42). *ρ* indicates Spearman’s correlation coefficients individually determined for mice treated with RNA-LNP or protein (protein group, *n*=6; RNA-LNP group, *n*=8).

We conclude that intravenous 1-18-LS RNA-LNP injection results in high antibody expression with a favorable pharmacokinetic profile and potent serum HIV-1 neutralizing activity that is maintained by repeated administrations.

### RNA-LNP-mediated expression of 1-18-LS RibobNAb durably suppresses viremia

To establish the antiviral activity of RNA-LNP-expressed 1-18-LS RibobNAb *in vivo*, we used CD34^+^ human hematopoietic stem cell-engrafted NXG mice with a high degree of humanization (average of 56.0% of hCD45^+^ cells amongst peripheral blood leukocytes) (**Figure S3A**). When effectively engrafted, CD34-humanized mice are susceptible to chronic HIV-1 infection, allow for the investigation of antiviral drug effects, and support the emergence of viral resistance in response to treatment-mediated selection pressure.^42^ After intraperitoneal challenge with recombinantly produced monoclonal HIV-1_YU2_, infection was confirmed by determining plasma viremia (**Figure S3B**). Four weeks after viral challenge, plasma viral loads ranged from 6,469 to 220,695 HIV-1 RNA copies/mL (geometric mean of 36,053 copies/mL) (**Figure 4A**). To ensure comparability between groups, mice were stratified into control and 1-18-LS RNA-LNP treatment arms based on baseline viral loads and stem cell donor distribution (**Figures S3C** and **S3D**).

**Figure 4.**
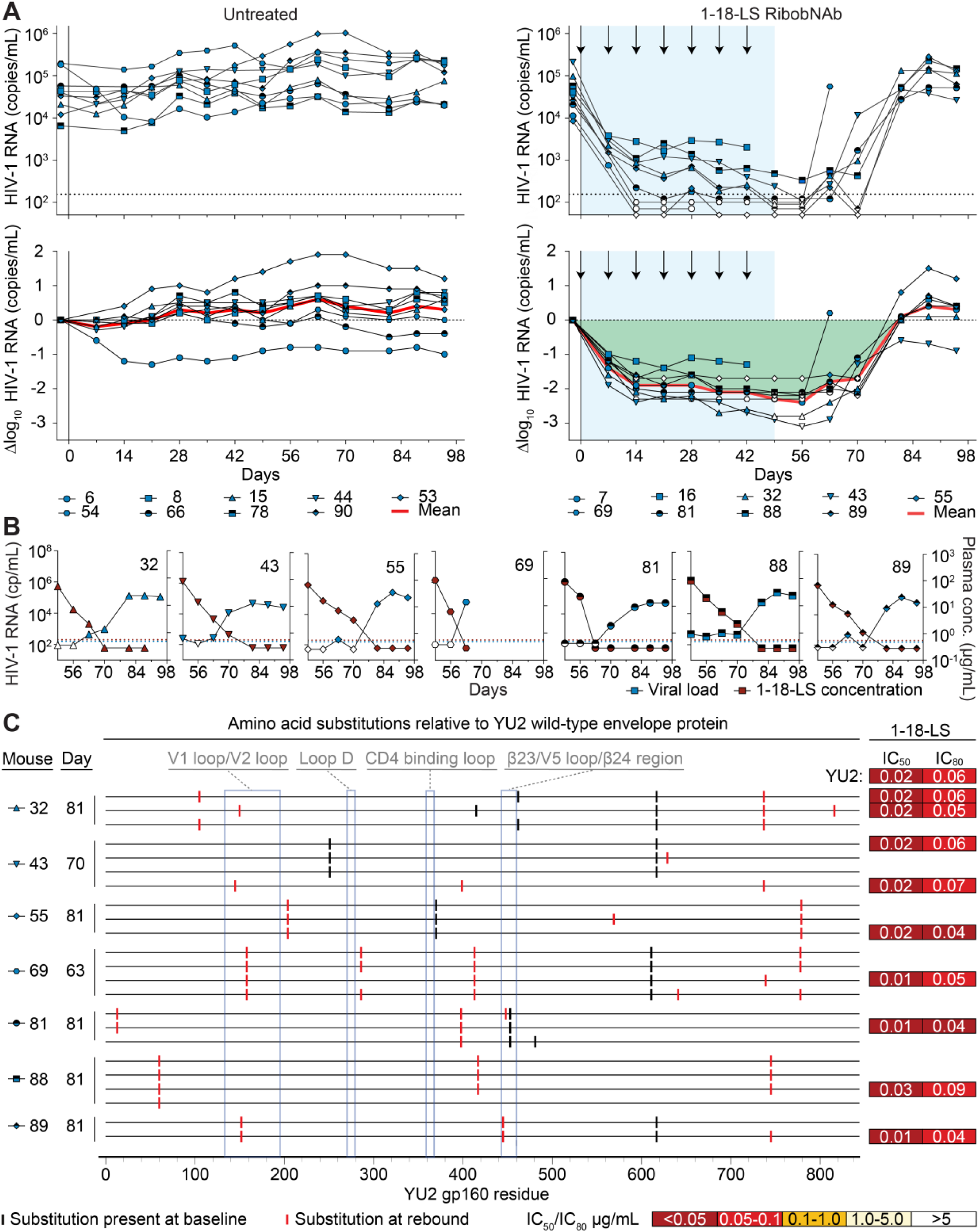
1-18-LS RibobNab monotherapy durably suppresses HIV-1. (**A**) HIV-1 RNA plasma copies (top) and log_10_ viral load changes compared with baseline (day -2) (bottom) in HIV-1_YU2_-infected CD34-humanized NXG mice. Dashed lines in top panels indicate HIV-1 RNA lower limit of quantification (LLOQ, 155 copies/mL) and white symbols indicate viral loads <LLOQ. Arrows indicate intravenous injections of 30 μg 1-18-LS RNA-LNP and total treatment period is indicated by blue shading. Red lines show average log_10_ viral load changes compared with baseline (*n*=9). (**B**) HIV-1 RNA plasma copies (left y axis, blue symbols) and plasma 1-18-LS RibobNAb levels (right y axis, red symbols) in individual mice indicated by IDs after termination of 1-18-LS RNA-LNP therapy. Blue and red dashed lines indicate HIV-1 RNA (155 copies/mL) and plasma 1-18-LS RibobNAb (0.56 µg/mL) LLOQs, respectively. White symbols indicate viral loads <LLOQ. (**C**) Plasma single genome amplification (SGA)-derived Env sequences obtained at viral rebound (days 63, 70, or 81). Amino acid substitutions relative to HIV-1_YU2_ wild-type Env are indicated in black if already present at baseline in the same mouse and in red if observed only at rebound. Boxes indicate IC_50_s and IC_80_s of 1-18-LS protein against YU2 wild-type pseudovirus and SGA-derived pseudoviruses; tested in technical duplicates.

In untreated control mice, overall stable viremia was observed throughout the 98-day observation period (**Figure 4A**). Weekly injections of LNP-formulated control RNA (encoding for incompatible antibody chains) did not reduce viremia, indicating that LNPs do not exhibit inherent antiviral activity (**Figure S3E**). In contrast, intravenous therapy with 30 µg 1-18-LS-encoding RNA-LNP rapidly reduced plasma viral loads in all treated mice (**Figure 4A**). Seven weekly RNA-LNP injections resulted in suppression of viremia to levels below the limit of quantification (LLOQ, 155 HIV-1 RNA copies/mL) in 7 out of 9 mice (77.8%) and an average maximum 2.4 log_10_ viral load drop compared to baseline (**Figure 4A**). Importantly, viremia remained suppressed compared to baseline in all NXG mice throughout the treatment period without viral rebound that would be suggestive of pre-existing or emerging resistance to 1-18-LS RibobNAb (**Figure 4A**). Average trough 1-18-LS RibobNAb plasma levels in RNA-LNP-treated mice ranged between 74 and 141 µg/ml and therapy resulted in potent plasma HIV-1 neutralization (**Figures S3F-S3H**).

After interruption of 1-18-LS RibobNAb treatment, all mice showed rebound of viremia within three to five weeks of the last RNA-LNP injection (**Figure 4A**). Rebound to near-baseline viral loads was associated with declining 1-18-LS RibobNAb plasma levels (**Figure 4B**). In all NXG mice studied, viral loads remained suppressed compared to baseline until 1-18-LS RibobNAb plasma concentrations dropped to less than 1 µg/mL, and 1-18-LS RibobNAb was mostly no longer detectable by anti-idiotype ELISA at the time of rebound (**Figure 4B**).

To determine potential selection for 1-18-LS RibobNAb-resistant viral variants during or after interruption of RNA-LNP treatment, we performed single genome amplification (SGA) of plasma HIV-1 *env* sequences at baseline and at rebound. Interspersed amino acid substitutions relative to the HIV-1_YU2_ wild-type sequence could be identified within Env before and/or after RNA-LNP treatment in all sequenced mice (**Figure 4C**). However, we did not find consistent selection of post-treatment mutations across multiple mice, including in the CD4 binding site targeted by 1-18-LS RibobNAb (**Figure 4C**). This suggests that the changes in HIV-1 Env observed after viral rebound were a consequence of random mutation rather than antibody-mediated selection. To confirm that the apparent lack of escape mutations after 1-18-LS RNA-LNP treatment was associated with a lack of viral resistance, we generated pseudoviruses derived from plasma *env* sequences and determined neutralization sensitivity to recombinant 1-18-LS protein. All tested post-rebound SGA-derived pseudoviruses remained highly sensitive to 1-18-LS within two-fold IC_50_ and IC_80_ ranges compared with HIV-1 YU2 wild-type pseudovirus (IC_50_ and IC_80_ ranges of 0.01 to 0.03 µg/mL and 0.04 to 0.09 µg/mL, respectively) (**Figure 4C**).

To compare the antiviral effects of RNA-encoded and recombinantly produced 1-18-LS, we treated HIV-1_YU2_-infected humanized NRG mice with weekly intravenous doses of 30 µg RNA-LNP or 500 µg protein. Indicating imperfect engraftment, mice showed a more limited degree of humanization (average of 10.1% hCD45+ cells) compared with NXG mice and hCD45+ cell reductions over time (5.1% vs 0.9% in mice tested before and at the conclusion of the experiment, *p*=0.008) (**Figures S2G** and **S2H**). While this is likely to have resulted in spontaneous reductions of viremia in some control mice and after interruption of treatment, treatment with RNA-LNP and recombinant protein showed comparable reductions of viremia compared with baseline during the first four weeks of therapy (**Figures S2I** and **S2J**).

Collectively, our results demonstrate that RNA-LNP-mediated 1-18-LS RibobNAb monotherapy can durably reduce and suppress viremia without emerging viral resistance.

### 1-18-LS RibobNAb maintains viral suppression of patient-derived viral isolates

Experiments with HIV-1_YU2_-infected humanized mice provide meaningful insights into viral escape pathways and can predict mutations emerging during bNAb therapy in humans.^42–45^ However, the use of recombinantly produced HIV-1_YU2_ that is partially based on a laboratory-adapted molecular clone does not fully recapitulate the genetic and phenotypic complexity of viruses circulating among people living with HIV. To determine the activity of RNA-LNP-formulated 1-18-LS RibobNAb against viruses that may be more challenging to control, we infected humanized mice with viral isolates derived from people living with HIV (**Figure 5A**; **Figures S4A** and **S4B**).^17^ After expansion of outgrowth culture viruses in MOLT-4 CCR5+ cells, we selected viruses originating from CD4 T cells of the ART-naïve individuals IDC#449 and IDC#548 for further analysis (**Figure 5A**). Based on plasma *pol* sequences, both viruses were classified as belonging to HIV-1 clade B. TZM-bl cell-based neutralization assays revealed susceptibility of bulk virus supernatants to recombinant 1-18-LS antibody, although sensitivity between the two viral swarms differed (IC_50_/IC_80_ values of 0.33/0.69 µg/mL and 0.04/0.10 µg/mL for IDC#449- and IDC#548-derived viruses, respectively) (**Figure 5B**).

**Figure 5.**
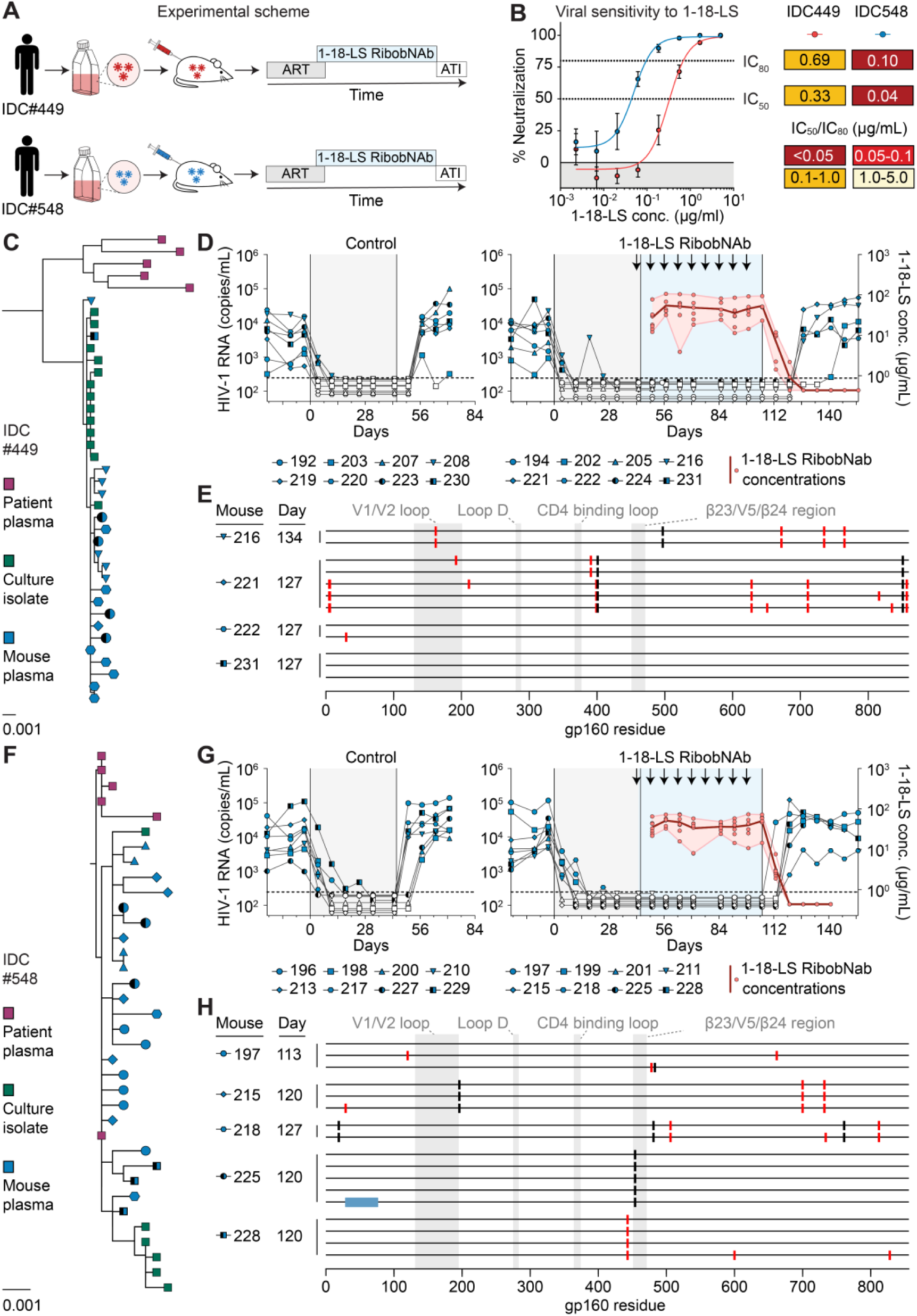
1-18-LS RibobNAb maintains suppression of patient-derived HIV-1 after ART interruption. (**A**) Experimental scheme. Bulk CD4 T cell outgrowth cultures were used to infect CD34-humanized mice subsequently treated with antiretroviral therapy (ART) for six weeks. RNA-LNP-treated mice received weekly intravenous injections of 30 μg 1-18-LS RNA-LNP starting 24 h prior to ART interruption. (**B**) Neutralizing activity of 1-18-LS recombinant protein against outgrowth virus culture supernatants used for challenges. Upper and lower dashed lines indicate 80% and 50% neutralization, respectively. Symbols and error bars indicate arithmetic mean and standard deviation of four independent assays performed in technical duplicates. (**C**) and (**F**) Phylogenetic trees of (C) IDC#449 and (F) IDC #548 HIV-1 *env* nucleotide sequences obtained by single genome amplification (SGA) from participant plasma, outgrowth culture supernatant, and mice at baseline (40 days after HIV-1 challenge). Bars indicate number of nucleotide substitutions per site. (**D**) and (**G**) Plasma HIV-1 RNA copies and 1-18-LS RibobNAb concentrations in mice infected with viral isolates from (D) IDC#449 and (G) IDC#548. Areas shaded in grey and blue indicate periods of ART and RNA-LNP treatments, respectively, and arrows indicate intravenous injections of 30 μg 1-18-LS RNA-LNP. Dashed lines indicate lower limits of quantification (LLOQ; HIV-1 RNA: 243 copies/mL; 1-18-LS RibobNAb: 0.88 µg/mL) and white symbols show viral loads <LLOQ (*n*=8 per group). Red lines indicate average 1-18-LS RibobNAb plasma concentrations, red symbols indicate individual measurements, and areas shaded in pink indicate concentration ranges. (**E**) and (**H**) Plasma SGA-derived Env sequences obtained at viral rebound. Bars indicate amino acid substitutions relative to mouse-specific baseline majority consensus sequence. Black bars indicate substitutions found in any baseline sequence of mice infected with the same viral isolate, red bars indicate substitutions only found in rebound sequences, and blue bars indicate amino acid deletions.

All mice challenged with IDC#548-derived virus (20/20) and 94% (17/18) of mice challenged with IDC#449-derived virus showed established HIV-1 infection three weeks after intraperitoneal exposure (**Figure S4C**). To determine potential viral selection or variation introduced during the viral outgrowth procedure or initial replication in infected mice, we performed SGA-derived *env* sequencing of RNA isolated from donor plasma, viral outgrowth cultures, and baseline mouse plasma (**Figures 5C** and **5F**; **Figures S5A** and **S5B**). Phylogenetic analysis revealed co-clustering of individual donor-derived *env* sequences obtained from all sample sources (**Figure S5A**). Comparison of infected mouse baseline consensus sequences demonstrated Env diversity between the viruses derived from IDC#449 and IDC#548 (76.2% and 72.9% amino acid identity for gp120/gp41 and gp120 only, respectively) (**Figure S5C**). Overall, *env* sequences isolated from infected mice resembled those identified in patient plasma and outgrowth cultures, and all studied mice showed some intra-host diversity in sequences identified at baseline (**Figures 5C** and **5F**; **Figures S5A** and **S5B**).

Results of early phase clinical trials indicate that HIV-1 bNAbs may contribute to long-acting therapies for maintaining control of viremia after initial daily antiretroviral therapy.^46–49^ To determine whether RNA-LNP-formulated 1-18-LS RibobNAb can effectively suppress patient-derived viruses in such a scenario, we first initiated ART in mice infected with IDC#449 or IDC#548 isolates (**Figure 5A**). After the start of oral antiretroviral treatment with a combination of tenofovir disoproxil fumarate, emtricitabine, and raltegravir, plasma viral loads dropped rapidly in 97% (36/37) of treated mice (**Figures 5D** and **5G**; **Figure S4D**). In most mice (94%), viral loads declined to levels below the qRT-PCR limit of quantification (243 HIV-1 RNA copies/mL) within two to three weeks of ART initiation (**Figures 5D** and **5G**; **Figure S4D**). Except for a small number of viral blips, viral loads in all suppressed mice remained below the limit of quantification throughout the ART treatment period (**Figures 5D** and **5G; Figure S4D**). Based on baseline viral loads and stem cell donor distribution, mice infected with each viral isolate were stratified into control and 1-18-LS RibobNAb treatment arms (**Figures S4E** and **S4F**). After completion of a six-week treatment period, ART was interrupted in all mice and plasma viral loads were continued to be determined weekly. Mice treated with 1-18-LS RibobNAb received weekly intravenous injections of 30 µg RNA-LNP starting one day prior to ART interruption (**Figure 5A**). Similar to observations made in people living with HIV that undergo treatment interruption, all control mice showed rapid viral rebound within one to two weeks of stopping antiretroviral therapy (**Figures 5D** and **5G**).^50^ In striking contrast, viremia continued to be fully suppressed in the absence of ART in all 1-18-LS RibobNAb-treated mice throughout the ten-week treatment period (**Figures 5D** and **5G**).

Indicating that control of viremia was mediated by 1-18-LS RibobNAb, interruption of RNA-LNP therapy resulted in viral rebound to near- or above-baseline levels within four weeks after the last injection in most mice (91%) (**Figures 5D** and **5G**). By the time of viral rebound, 1-18-LS RibobNAb plasma levels had declined below the quantification limit of the antibody concentration assay (<0.88 µg/mL) in 91% (10/11) of studied mice (**Figures 5D** and **5G**; **Figure S4G**). In the single mouse with quantifiable 1-18-LS RibobNAb at rebound (mouse 197), antibody levels seven days after the last injection were lower than in all other tested mice (14.4 µg/mL vs. average of 55.7 µg/mL [range: 23.0-93.7]) and had declined to only 22.4-fold (2.24 µg/mL) above the 1-18-LS IC_80_ against the viral challenge stock at rebound (**Figure S4G**). Compared with baseline, plasma *env* sequences obtained after interruption of 1-18-LS RibobNAb treatment and rebound showed limited changes (**Figures 5E** and **5H**). Detected post-rebound-specific amino acid substitutions were restricted to regions outside of the CD4 binding site and we observed no consistent selection for recurring amino acid substitutions across the different animals (**Figures 5E** and **5H**). Collectively, these results suggest that no resistance to 1-18-LS RibobNAb emerged for both of the tested patient isolates over more than eight weeks of exposure.

We conclude that RNA-LNP-mediated expression of 1-18-LS RibobNAb is effective in maintaining ART-free suppression of clinically relevant patient-derived HIV-1 isolates.

## DISCUSSION

Early phase clinical trials of broadly neutralizing antibodies demonstrated the potential of antibody-mediated strategies to treat and prevent HIV-1 infection.^51^ Despite promising developments, vaccines capable of inducing HIV-1 bNAbs remain elusive and vector-mediated expression strategies such as the use of adeno-associated viruses did not yet achieve meaningful bNAb serum levels in humans.^52–54^ Thus, current HIV-1 bNAb-based approaches rely on the parenteral administration of recombinantly produced monoclonal antibodies.^55^ While half-life optimization facilitates long dosing intervals (e.g., every three to six months), clinical application of bNAbs for HIV-1 suppression will likely require life-long therapy. Although LNP-formulated RNA also requires specialized processing and appropriate storage conditions, its application could address some of the challenges and costs of protein-based drugs.^3,4^ Using LNP-formulated RNA encoding the potent 1-18-LS RibobNAb, we demonstrate effective immunotherapy of HIV-1 infection based on endogenous bNAb expression that is independent of recombinant antibody manufacturing and administration.

Intravenous administration of RNA-LNPs typically results in efficient delivery to the liver. Optimized LNP formulations combined with nucleoside-modified RNA can minimize LNP-induced inflammatory responses and RNA sensing by innate immune receptors to enhance RNA translation.^56–58^ Consistent with RNA-mediated bNAb expression kinetics, 1-18-LS RibobNAb concentrations peaked over the first 2-3 days of i.v. RNA-LNP injection and showed a limited decline within the first week. Altogether, RNA-expressed 1-18-LS RibobNAb showed a favorable pharmacokinetic profile in diverse mouse models, including an hFcRn-transgenic model with established predictive value for human antibody PKs.^40^ Studies in non-human primates (NHPs) demonstrated comparable serum half-lives of an HIV-1 antibody administered as recombinant protein or expressed from intravenously administered RNA-LNP constructs.^59^ Moreover, in a phase 1 clinical trial, an LS-modified RNA-encoded CHIKV antibody showed a long half-life (≈69 days) after RNA-LNP infusion.^60^ Based on the human serum half-life of 1-18-LS predicted by scaling from NHP and mouse studies (≈50 days),^37^ these observations and our results suggest that 1-18-LS RibobNAb can be developed as a long-acting strategy with administration intervals similar to recombinant bNAbs (i.e., every three to six months), if sufficient antibody levels can be achieved.

We observed high 1-18-LS RibobNAb levels up to >1,000 µg/mL after a 30 µg RNA-LNP injection in mice (i.e., ca. 1,500 µg/kg body weight). While 1-18-LS RibobNAb expression in humans remains to be determined, a 600 µg/kg dose of RNA encoding for chikungunya antibody CHKV-24 resulted in only moderate peak serum antibody levels of ≈10 µg/mL in humans.^60^ In addition to using optimized LNP formulations and RNA for enhanced antibody expression in humans, selection of potent bNAbs to maximize antiviral activity at a given antibody level will therefore be critical for RNA-based strategies targeting HIV-1. Given their broad activity, CD4 binding site antibodies are likely to serve as a backbone of antibody treatment regimens and, accordingly, represent the most extensively studied bNAb class.^25–30,32,46–48,51,61–67^ Monotherapy with clinically advanced CD4bs bNAbs in the relevant HIV-1_YU2_-infected mouse model typically results in rapid viral escape.^17,22^ In contrast, and similar to the results for recombinant 1-18 antibody given at total weekly subcutaneous doses of 1,000 µg of protein,^17^ 1-18-LS RibobNAb therapy with weekly i.v. injections of 30 µg RNA-LNP effectively suppressed HIV-1_YU2_ viremia and prevented viral escape. Moreover, maintained suppression of infection with diverse patient-derived viral isolates in the absence of ART emphasizes the broadly potent activity of 1-18-LS RibobNAb and provides first *in vivo* evidence for 1-18-LS activity against non-recombinant viruses.

In humanized mice, 1-18-LS RibobNAb monotherapy demonstrated excellent efficacy in reducing viral loads and preventing HIV-1 rebound. Nevertheless, low levels of viral RNA remained detectable in some mice with adequate antibody expression. Following a rapid and substantial drop in viral loads after treatment initiation, viremia showed a more gradual decline over time in most of these mice. Although the reason for the lack of rapid suppression to entirely undetectable viral RNA in some mice remains unclear, it may be related to the bNAb mechanism of action and is not uncommon for effective bNAb therapy in viremic mice with high baseline viral loads.^17,22,45,68^ Irrespective of their use in viremic or pre-treated individuals, clinical application of antibody-based strategies will likely require co-administration of antiretroviral drugs and/or antibody combinations to minimize the potential of escape and resistance. Co-formulation of different coding RNA chains into single lipid nanoparticles facilitates co-expression of different proteins in single transfected cells.^69^ However, combining heavy and light chain-coding RNAs of different antibodies into single or different LNPs carries the risk for heavy/light chain mispairings in co-transfected cells that may result in reduced expression, increased immunogenicity, and/or off-target reactivity.^70^ Thus, future RNA-based approaches for expression of bNAb combinations will require antibody engineering strategies that effectively prevent mispairing while maintaining neutralizing potency of parental antibodies.

Collectively, our results provide a proof-of-concept for effective RNA-encoded bNAb immunotherapy of HIV-1 infection and support the development of RNA-based antibody therapeutics for viral infections.

## MATERIALS AND METHODS

### Mouse models

NOD.Cg-*Rag^1tm1mom^Il2rg^tm1Wjl^*/SzJ (NRG) mice and NOD.Cg-*Fcgrt^tm1Dcr^ Prkdc^scid^ Il2rg^tm1Wjl^* Tg(FCGRT)32Dcr/J (NSG hFcRn Tg32) mice were purchased from The Jackson Laboratory; CD34-humanized NOD-*Prkdc^scid^-IL2rg^Tm1^*/Rj (NXG) mice were purchased from Janvier Labs. NRG mice were bred under specific pathogen-free conditions with 12-hour light/dark cycles. CD34-humanized NRG mice were generated as previously described.^17,45^ In brief, human CD34^+^ hematopoietic stem cells were isolated from cord blood using CD34 microbeads (Miltenyi Biotec). Within 5 days of birth, mice were sublethally irradiated and injected intrahepatically with human CD34^+^ cells 4 to 6 hours after irradiation. Humanization was confirmed by flow cytometry as indicated below. For experiments involving antiretroviral therapy, NXG mice were provided ssniff 1154 breeding feed with added antiretroviral drugs (see below). All experiments involving NRG and NXG mice were authorized by the State Agency for Consumer Protection and Food North Rhine-Westphalia (LANE NRW), and experiments involving NSG hFcRn Tg32 mice were approved by the regional animal welfare committee. All mouse studies were conducted in accordance with FELASA recommendations, the German Animal Welfare Act, and EU Directive 2010/63/EU.

### Clinical samples

Cord blood for humanization of NRG mice was obtained with written informed consent under protocols approved by the Ethics Committee of the Medical Faculty of the University of Cologne (16-110, 18-420) and the Ethics Committee of the North Rhine State Chamber of Physicians (2018382). Samples from people living with HIV were collected with written informed consent under protocols approved by the Ethics Committee of the Medical Faculty of the University of Cologne (13-364, 16-054).

### Humanization analysis of CD34-humanized mice

Engraftment of CD34-humanized NRG mice was assessed by flow cytometry on an Aria Fusion (BD). EDTA blood samples were stained using anti-human antibodies for CD19 (APC), CD8 (FITC), CD3 (Pacific Blue), CD4 (PE; all from BD), and CD45 (Pacific Orange, Thermo Fisher), as well as using an anti-mouse CD45 antibody (PE/Cy7, BioLegend) for 20 min at 4°C in FACS buffer (PBS containing 2% FBS and 2 mM EDTA). Erythrocytes were then lysed using ACK lysis buffer (Thermo Fisher), and samples were washed and analyzed in FACS buffer. To determine humanization parameters of HIV-1_YU2-_infected NRG mice, cells were fixed after the erythrocyte lysis step using 2% formaldehyde (Thermo Fisher) for 15 min at room temperature. Analysis was performed using FlowJo (BD). For CD34-humanized NXG mice, humanization data and data regarding stem cell donors were provided by Janvier Labs.

### Cell lines

HEK293T cells were maintained at 37°C and 5% CO_2_ in Dulbecco’s Modified Eagle Medium (DMEM, Thermo Fisher) supplemented with 10% fetal bovine serum (FBS, Sigma), 1 mM sodium pyruvate, 100 U/mL penicillin, 100 µg/mL streptomycin, 0.25 µg/mL amphotericin B, and 2 mM of L-Glutamine (all Thermo Fisher). For RNA transfection experiments, HEK293T/17 cells (LGC Limited) were cultured in DMEM medium supplemented with GlutaMAX (Thermo Fisher) and 10% FBS (Thermo Fisher) and maintained at 37°C and 7.5% CO_2_ in a humidified incubator. TZM-bl cells were maintained at 37°C and 5% CO_2_ in DMEM supplemented with 10% FBS, 50 µg/mL gentamicin (Sigma), 1 mM sodium pyruvate, 2 mM L-glutamine, and 25 mM HEPES (Sigma). MOLT-4 CCR5+ cells were maintained at 37°C and 5% CO_2_ in RPMI 1640 medium (Thermo Fisher) supplemented with 10% FBS, 100 U/mL penicillin, 100 µg/mL streptomycin, and 1 mg/mL of geneticin (G-418) (Thermo Fisher). The sex of HEK293T, HEK293T/17, and TZM-bl cell lines is female. The sex of MOLT-4 CCR5+ cells is male.

### RNA and RNA-LNP production

1-18-LS-encoding RNAs were produced as previously described via *in vitro* transcription of codon optimized DNA sequences to generate heavy and light chain-encoding RNAs.^7^ Variable domain sequences of 1-18 heavy and light chains were cloned into the multiple-cloning site (MCS) of pST1-hAg-Kozak-MCS-F-I-A30LA70 that contains human kappa light chain or IgG1 heavy chain (with or without LS substitutions) constant domains (modified from ref. ^71^). RNA transcripts contain conserved structural elements promoting RNA stability and efficient protein translation. These regulatory elements consist of a 5’-untranslated region (AGA-dEarI-hAg), a CC413 cap at the 5’ end, a 3’-untranslated region containing the FI element and a 100-nucleotide poly-adenine tail with an integrated linker at the 30^th^ position (A30L70). RibobNAb-encoding DNA sequences were transcribed *in vitro* using T7 RNA polymerase (Ambion), CleanCap 413 (TriLink Biotechnologies) for 5′ capping, and modified nucleotide triphosphates, producing N1-methylpseudouridine (m1Ψ)-capped RNA. RNA products were isolated using magnetic bead separation, followed by cellulose purification and resuspension in ribonuclease-free water (B. Braun) containing 10 mM HEPES and 0.1 mM EDTA (BioChrom). Concentrations of synthesized RNAs were determined by spectrophotometry (NanoDrop 2000c, Thermo Fisher) and quality control was performed using capillary electrophoresis (Agilent 2100 Bioanalyzer, Agilent Technologies). RNAs were stored in nuclease-free tubes (Eppendorf) at −65°C to −85°C and either used for formulation into lipid nanoparticles (LNPs) or for RibobNAb expression using RiboJuice or electroporation. To generate 1-18-LS RNA-LNP variants, the respective heavy- and light-chain encoding 1-18-LS RNAs were mixed in a 1.5:1.0 weight ratio and encapsulated in an LNP formulation consisting of an ionizable cationic lipid, a polyethylene glycol (PEG) lipid, phospholipid, and cholesterol, followed by storage at −65°C to −85°C. Control RNA-LNPs were generated by combining heavy and light chain sequences of two different antibodies.

### Production of recombinant antibody

Recombinant 1-18 and 1-18-LS protein was produced by LakePharma by transient transfection of CHO cells, followed by purification using protein A chromatography and size exclusion chromatography. Purified antibody was stored at −65°C to −85°C.

### *In vitro* RibobNAb expression

To express RibobNAbs for binding experiments, HEK293T/17 (passaged at least three times) were seeded at a density of 4x10^5^ cells/well in a 12-well plate (Greiner Bio-One) and cultured for 24 h at 37°C and 7.5% CO_2_. 1 µg total RNA was transfected per well at a heavy chain:light chain RNA ratio of 1.5:1.0 (0.6 µg and 0.4 µg). RiboJuice^TM^ mRNA transfection reagents were prepared according to the manufacturer’s instructions (Merck Millipore). Complexes with RNA were added dropwise to the cells and mixed by gently shaking the plate. Subsequently, cells were incubated at 37°C and 7.5% CO_2_ in a humidified incubator for 48 h. Cell culture supernatants were centrifuged at 400 x *g* for 2 min, followed by another centrifugation at 3,000 x *g* for 10 min to remove any residual cells or debris. Cell culture supernatants were stored at 4°C until analysis and antibody concentration in supernatants was determined by Gyros ELISA.

To express RibobNAbs for neutralization assays, HEK293T/17 cells were seeded at a density of 1x10^4^ cells/cm^2^ in a T175 suspension flask. After 24h, cells were centrifuged at 300 x *g* for 8 min and washed twice with chilled X-VIVO 15 medium (Lonza). 2x10^6^ cells at a density of 8x10^6^ cells/mL in X-VIVO 15 medium were transferred into pre-cooled 0.4 cm electroporation cuvettes and 25 µg RNA (15 µg heavy chain RNA, 10 µg light chain RNA) was electroporated at 250 V using two 5 ms pulses with a 400 ms interval. After incubation on ice for 10 min, cells were resuspended in 750 µL Expi293™ medium (Thermo Fisher), seeded into a 12-well plate at a density of 2x10^6^ cells/well, and incubated at 37°C and 5% CO2 in a humidified incubator for 48 h. Subsequently, cell culture supernatants were centrifuged at 300 x *g* for 10 min and stored at 4°C until analysis. Antibody concentration in supernatants was determined by Gyros ELISA.

### RNA-LNP lipofection

HEK293T/17 cells were passaged at least three times prior to lipofection. Cells were seeded at a density of 4x10^5^ cells/well in a 12-well plate (Greiner Bio-One) 24 h before lipofection and incubated overnight at 37°C with 7.5% CO_2_. Subsequently, cell culture supernatants were aspirated and replaced with 800 µL prewarmed Opti-MEM-I reduced serum medium (Thermo Fisher). RNA-LNP aliquots were thawed and thoroughly mixed by gently flipping the tube without vortexing. RNA-LNPs (1 µg RNA/well) were prepared in 200 µL Opti-MEM-I reduced serum medium and added dropwise. Plates were gently swirled to evenly distribute RNA-LNP onto the cells, followed by incubation at 37°C and 7.5% CO_2_ in a humidified incubator for 48 h. At the end of the incubation period, cell culture supernatants were centrifuged at 400 x *g* for 2 min, followed by another centrifugation at 3,000 x *g* for 10 min to remove any residual cells or debris. Cell culture supernatants were stored at 4°C until analysis and antibody concentration in supernatants was determined by Gyros ELISA.

### SDS PAGE and Western blot

Cell culture supernatants of HEK293T/17 cells transfected with 1-18- or 1-18-LS-encoding RNA or lipofected with 1-18-LS RNA-LNP, and NSG hFcRn Tg32 mouse serum samples collected 24 hours after RibobNAb injection were diluted in DPBS and mixed with 4x Laemmli buffer. For reducing conditions, dithiothreitol (Carl Roth) was added to a final concentration of 100 mM. Samples were denatured at 95°C for 5 min. A total of 7 ng of protein for cell culture supernatants and 10 ng for mouse serum samples were loaded onto precast 4-15% Criterion™ gradient SDS-PAGE gels (Bio-Rad) and separated for 40 min at 250 V and 2-8°C in TGS running buffer (Bio-Rad). 7 ng of recombinantly produced and purified 1-18 or 1-18-LS was included as reference in addition to10 µL of 1:1 mixed Precision Plus Protein^TM^ All Blue Prestained and Unstained Protein Standards (Bio-Rad) for supernatant samples and 10 µL of Precision Plus Protein^TM^ Dual Color Standards (Bio-Rad) for serum samples. For western blotting, proteins were transferred onto a Trans-Blot Turbo Midi 0.2 µm nitrocellulose membrane (Bio-Rad) at 2.5 A for 7 min using the Trans-Blot® Turbo^TM^ blotting system (Bio-Rad). Membranes were blocked for 1 h at room temperature with 5% skimmed milk dissolved in Tris-buffered saline (TBS) containing 0.5% Tween-20 (TBST 0.5%, Sigma). After blocking, membranes were washed with TBS containing 0.1% Tween-20 (TBST 0.1%) and incubated with a Peroxidase-conjugated goat anti-human-Fcγ-specific antibody (Jackson ImmunoResearch) diluted 1:1,000 and a peroxidase-conjugated goat anti-human kappa light chain antibody (Biozol) diluted 1:500 in TBST 0.5% supplemented with 3% BSA Fraction V (Eurobio). For mouse serum samples, membranes were stained with Peroxidase-conjugated goat anti-human-Fcγ-specific detection antibody (Jackson ImmunoResearch) diluted 1:500 and a peroxidase-conjugated goat anti-human kappa light chain detection antibody (Thermo Fisher) diluted 1:200. Membranes were incubated for 1 h and washed with TBST 0.1%. Protein bands were visualized with Clarity^TM^ Western ECL Reagent (Bio-Rad) on a Fusion FX imaging device (Vilber) with an exposure time of 0.1 s for cell culture supernatant samples and 0.5 s for mouse serum samples and merged with white light image to visualize the marker. Data analysis was performed with the Image Lab Software (Bio-Rad).

### SOSIP ELISA

HIV-1 Env binding of recombinant 1-18-LS protein and RibobNAb 1-18-LS in cell culture supernatants was evaluated by ELISA using 96-well streptavidin plates (Thermo Scientific). Wells were coated with 100 ng biotinylated recombinant BG505 DS-SOSIP.664 (ATUM; custom-made) in 100 µL coating buffer (16 mM sodium carbonate and 34 mM sodium hydrogen carbonate; pH 9.6) and incubated overnight at 4°C. Blocker™ Casein in PBS (Thermo Fisher) was diluted to 0.18% casein (w/v) with PBS and used as blocking buffer. Plates were washed three times with PBS-T (phosphate-buffered saline containing 0.01% Tween-20) and blocked with 250 µL blocking buffer per well for 1 h at 37°C with gentle shaking. After another round of washing, serially diluted recombinant 1-18-LS in blocking buffer and cell culture supernatants as well as negative control IgG were added in duplicates and incubated at 37°C for 1 h on a shaker. After washing, horseradish peroxidase-conjugated goat anti-human IgG antibody was added (1:5,000 in blocking buffer; Jackson ImmunoResearch), and plates were incubated at 37°C for 45 min on a shaker. Plates were washed again, and 100 µL of 3,3′,5,5′-tetramethylbenzidine substrate (Kementec) was added and incubated for 8 min at room temperature in the dark. The reaction was stopped with 100 μL 25% sulfuric acid, absorbance at 450 nm and 620 nm was measured on an Epoch microplate reader (BioTek), and mean differences in optical density (ΔOD450-620 nm) were calculated.

### Biolayer interferometry

Biolayer interferometry was performed using the Octet®-RH96 Protein Analysis System (Sartorius). All experiments were performed at 30°C with orbital agitation at 1,000 rpm. To measure affinity of 1-18-LS recombinant protein and 1-18-LS RibobNAb to a stabilized HIV-1 Env trimer, antibodies or cell culture supernatants were immobilized as ligands on the surface of Octet® anti-hIgG Fc Capture (AHC) Biosensors (Sartorius), while soluble BG505 DS-SOSIP.664 (NIH) was used as analyte. Prior to antibody immobilization, a baseline step using Octet® Kinetics Buffer (Sartorius) was performed for 60 s. Antibodies were immobilized at a concentration of 1 µg/mL for 900 s with a threshold of 1 nm, followed by a baseline step in Octet® Kinetics Buffer for 120 s. Binding affinities were determined using multi-cycle kinetics and decreasing BG505 DS-SOSIP.664 concentrations (26.71 nM to 0.42 nM; 2-fold dilutions in Octet® Kinetics Buffer). The association step was carried out for 600 s, followed by a 600 s dissociation step in Octet® Kinetics Buffer. Between cycles, sensors were regenerated five times by incubating sensors in glycine buffer (pH 1.5) for 5 s, followed by a 5 s neutralization in Octet® Kinetics Buffer. Data were analyzed using the Octet® Analysis Studio 13.0.1.35 software. To perform inter-step correction, data were aligned with the dissociation step, and Savitzky-Golay filtering was applied to all curves. Association and dissociation curves were globally analyzed using a 1:1 Langmuir binding model.

### Sample collection for serum/plasma antibody levels and neutralization titers

For pharmacokinetic profile analysis in hFcRn-transgenic mice, 30 µg or 10 µg 1-18-LS RNA-LNP, or 100 µg recombinant 1-18-LS protein (each diluted in DPBS to a final volume of 150 µL) was administered as a single i.v. injection. Mice were randomly distributed to treatment groups and each group was divided into two subgroups for longitudinal rotational sampling. Blood was collected using Microvette 500 Z-Gel tubes (Sarstedt), centrifuged at 10,000 x *g* for 5 min, and serum was stored at -65°C to -85°C until analysis by Gyros ELISA.

For pharmacokinetic profile analysis in HIV-1-negative non-humanized NRG mice, 30 µg 1-18-LS RNA-LNP or 200 µg recombinant 1-18-LS protein (each diluted in PBS to a final volume of 100 µL) were administered as a single i.v. injection. Serum was obtained by centrifugation of serum collection tubes (Sarstedt) at 10,000 x *g* for 5 min, and samples were stored at -20°C before analysis by Gyros ELISA or in pseudovirus neutralization assays as indicated below.

For the analysis of trough antibody concentrations and neutralizing activity in HIV-1_YU2_-infected CD34-humanized NRG mice, blood samples were collected every seven days prior to i.v. injection of 30 µg 1-18-LS RNA-LNP or 500 µg recombinant 1-18-LS protein (each diluted in DPBS (Thermo Fisher) to a final volume of 150 µL). Serum was obtained by centrifugation of serum collection tubes at 10,000 x *g* for 5 min and samples were stored at -20°C before analysis by Gyros ELISA or in pseudovirus neutralization assays as indicated below. Samples for ELISA were inactivated after a 2.22-fold dilution in DPBS containing Triton X-100 (Carl Roth) at a final concentration of 1%.

For the analysis of antibody concentrations and neutralizing activity in HIV-1_YU2_-infected CD34-humanized NXG mice, EDTA plasma samples were collected prior to weekly i.v. injections of 30 µg 1-18-LS RNA-LNP diluted in DPBS to a final volume of 150 µL or no treatment. EDTA plasma was obtained by centrifugation of EDTA collection tubes (Sarstedt) at 2,000 x *g* for 10 min and stored at -20°C before analysis by Gyros ELISA or in pseudovirus neutralization assays as indicated below. Samples for ELISA were inactivated after a 2.22-fold dilution in DPBS containing Triton X-100 at a final concentration of 1% (except for a single sample diluted 4.44-fold and five samples diluted 12.21-fold).

For the analysis of antibody concentrations in NXG mice infected with patient isolates, EDTA samples were collected prior to i.v. injections of 30 µg 1-18-LS RNA-LNP or weekly thereafter, and plasma samples were collected and inactivated as indicated above after a 2.22- or 11.1-fold dilution in DPBS.

### Recombinant HIV-1 production

Replication-competent HIV-1_YU2_ (HIV-1 YU2 Env in the HIV-1 NL4-3 lab strain backbone) was produced by transfection of HEK293T cells with an infectious molecular clone using FuGENE 6 (Promega).^72^ Cell culture supernatants containing replication-competent virus were harvested after 24 h and 48 h, and stored at -80°C before mouse challenge.

### ELISA for antibody quantification

RibobNAb levels in cell culture supernatants and mouse serum or plasma were quantified by Gyrolab® automated immunoassay using kits, microtiter assay plates, measuring device, and analysis software from Gyros Protein Technologies AB. For quantification of RibobNAbs in cell culture supernatant, Reagent A (Gyrolab® huIgG Kit), which contains a biotinylated protein A derivative, or 0.1 mg/mL biontinylated anti-human Fc-specific antibody (Thermo Fisher) was used for capture. For quantification of RibobNAbs or recombinant 1-18 in mouse serum or plasma, Reagent A (Gyrolab® Generic PK or TK Kit), which contains a biotinylated anti-human IgG Fc-specific antibody, or 0.1 mg/mL of custom-made biotinylated anti-1-18 F(ab’)₂ antibody (BioRad) was used for capture. Detection was performed using Reagent B from the same kits (Gyrolab® huIgG Kit, Gyrolab® Generic PK or TK Kit), which contain an Alexa Fluor 647-labeled F(ab’)2 fragment of anti-human IgG, or 25 nM of an Alexa Fluor-647 labeled F(ab’)₂ fragment rabbit anti-human IgG Fcy (Jackson ImmunoResearch). Capture and detection antibodies were diluted in PBS/Tween 20 buffer (Thermo Fisher) and Rexxip^TM^ F buffer (Gyros Protein Technologies), respectively. Assays were processed in a Gyrolab® Bioaffy 1000 HC CD (Generic PK Kit, huIgG Kit) or Gyrolab® Bioaffy 20 HC CD (Generic TK kit). Cell culture samples were diluted in Reagent E buffer (Gyrolab® huIgG Kit) at 10-fold for RiboJuice-transfected and RNA-LNP-lipofected samples. Serum or plasma samples (in some cases after inactivation and predilution) were diluted 10-fold, 20-fold, or 100-fold with Reagent F buffer (from Gyrolab® Generic PK or TK Kit). The CD columns were washed with Reagent C and Reagent D (Gyrolab® huIgG Kit). An eleven-point three-fold serial dilution standard curve was generated using an IgG1 reference protein (Gyrolab® Gyros Protein Technologies, LakePharma) or recombinant 1-18-LS (LakePharma) diluted in Reagent E buffer for cell culture supernatant and Reagent F buffer for serum samples. Dilutions were made in buffers spiked with sample-matched matrix at the final dilution factor. Data were generated on a Gyrolab® xPand^TM^ ELISA device (Gyros Protein Technologies) and analyzed with Gyrolab® Evaluator software.

### HIV-1 infection and treatment of HIV-1-infected mice

CD34-humanized NRG or NXG mice were challenged intraperitoneally twice within 2-3 days. Mice were treated weekly by i.v. injection of 30 µg of 1-18-LS RNA-LNP or 500 µg 1-18-LS recombinant protein diluted in DPBS (Thermo Fisher) or DPBS only at a final volume of 150 µL. Mice in untreated control groups received no injections. Oral antiretroviral therapy was provided through ad-libitum access to food pellets containing tenofovir disoproxil (provided as tenofovir disoproxil fumarate, Zentiva), emtricitabine (Zentiva), and raltegavir (MSD) at doses of 720, 586.76, and 4,800 mg per kg of food weight (ssniff Spezialdiäten GmbH).

### Viral load measurements

EDTA plasma was collected after centrifugation of EDTA blood collection tubes (Sarstedt) at 2,000 x *g* for 10 min. RNA was extracted using the QIAamp MinElute Virus Spin Kit (Qiagen), including a DNase I (Qiagen) digestion step, on a Qiacube device (Qiagen). In HIV-1_YU2_-infected mice, viral loads were determined by quantitative real-time qPCR (qRT-PCR) using *pol*-specific primers 5′-TAATGGCAGCAATTTCACCA and 5′-GAATGCCAAATTCCTGCTTGA as well as internal probe 5′-/56-FAM/CCCACCAACARGCRGCCTTAACTG/ZenDQ/ (all IDT) ^68^. In mice infected with primary isolates, LTR-specific primers 5′-CTCAATAAAGCTTGCCTTGA and 5′-GGGCGCCACTGCTAGAGA as well as internal probe 5′-/56-FAM/AGTAGTGTGTGCCCGTCTGT/ZenDQ/ (all IDT) were used.^73^ qRT-PCR was performed using the Taqman RNA-to-CT 1-step kit (Thermo Fisher) on a QuantStudio cycler (Thermo Fisher). qRT-PCR was run for 15 min at 48°C and 10 min at 95°C, followed by 45 cycles of 15 s at 95°C and 1 min at 60°C. For quantification, an HIV-1_YU2_ standard was produced by infection of SupT1-R5 cells with HIV-1_YU2_, heat-inactivation of cell culture supernatant, and dilution in Tris-buffered saline. HIV-1 RNA copy number of the standard was assessed using the quantitative Alinity m HIV-1 kit (Abbott) or the quantitative cobas 6800 HIV-1 kit (Roche). The lower limit of quantification was determined as 155, 243, or 387 HIV-1 RNA copies/mL, depending on the standard production batch. A single standard batch was used throughout each independent experiment. Standards (1:5 dilution series in AVE buffer, Qiagen) were included for every qRT-PCR run and viral loads were calculated using the QuantStudio software.

### Single genome amplification and sequencing

Single genome amplification and sequencing of HIV-1 *env* were performed as previously described.^17,74^ Briefly, viral RNA was extracted from EDTA plasma or culture supernatant using the QIAamp MinElute Virus Spin Kit (Qiagen), including a DNase I (Qiagen) digestion step, on a Qiacube device (Qiagen). cDNA was generated using Superscript IV (Thermo Fisher) and antisense primer YB383 5′-TTTTTTTTTTTTTTTTTTTTTTTTRAAGCAC followed by an RNase H (Thermo Fisher) digestion step at 37°C for 20 min. *env* cDNA was then amplified by nested PCR with limiting dilutions. The first round of PCR amplification was performed using primers YB383 5′-TTTTTTTTTTTTTTTTTTTTTTTTRAAGCAC and YB50 5′-GGCTTAGGCATCTCCTATGGCAGGA AGAA. First PCR was performed at 98°C for 45 s, followed by 35 cycles of 98°C for 15 s, 55°C for 30 s, and 72°C for 15 min, and a final extension at 72°C for 15 min. For the second PCR, 1 µL of first PCR product was used as template and PCR was performed using primers YB49 5′-TAGAAAGAGCAGAAGACAGTGGCAATGA and YB52 5′-GGTGTGTAGTTCTGCCAATCAGGGAAGWAGCCTTGTG. Second PCR was performed at 98°C for 45 s, followed by 45 cycles of 98°C for 15 s, 55°C for 30 s, and 72°C for 15 min, and a final extension at 72°C for 15 min. PCRs were performed using Phusion Hot Start Flex DNA Polymerase (New England Biolabs), and second PCR products were sequenced by Sanger sequencing or Oxford Nanopore Technologies long-read next-generation sequencing. Typically, only dilutions leading to <30% positive PCRs were further analyzed, so that >80% of positive PCRs are amplified from a single virion (according to Poisson’s distribution). In a limited number of cases, cDNA dilutions resulting in up to a maximum of 33% positive reactions were also analyzed.

### Env cloning

SGA-derived HIV-1 *env* sequences were cloned into expression vectors for pseudovirus production. A vector backbone was generated by PCR-mediated linearization of the pcDNA.3.1D V5-His-TOPO plasmid (Thermo Fisher) with the Q5 Hot Start High-Fidelity DNA polymerase (New England Biolabs) using primers pcDNA3.1_Xho1 5′-GATCCGAGCTCGGTACCAAGC and pcDNA3.1_BamH1 5′-TCGAGTCTAGAGGGCCCGCGG. For *env* amplification, first PCR SGA products were used as templates for PCR using Phusion Hot Start Flex DNA Polymerase (New England Biolabs) with the primers HIV_EnvB3in_SLIC 5′-CCGCGGGCCCTCTAGACTCGAGTCTCGAGATACTGCTCCCACCC and HIV_EnvB5in_SLIC 5′-GCTTGGTACCGAGCTCGGATCTTAGGCATCTCCTATGGCAGGAAGAAG with overhangs matching the linearized backbone. Both linearized vector and *env P*CR products were gel-purified using the NucleoSpin Gel and PCR Clean-up kit (Macherey-Nagel). For assembly of linearized backbone plasmid and *env* PCR product into a single expression plasmid, sequence and ligation independent cloning (SLIC) was performed using T4 DNA polymerase (New England Biolabs). Briefly, 50 ng of linearized backbone and 100 ng of *env* PCR product were mixed in 10 µL H₂O with 1 µL buffer r2.1. T4 DNA polymerase (0.2 µL) was added and incubated at room temperature for exactly 4 min, after which 1 µL of 10 mM dGTP (Thermo Scientific) was added to limit exonuclease activity. The reaction was then heat-inactivated at 65°C for 5 min and transformed into DH5a or STBL3 cells. Alternatively, the NEBuilder HiFi DNA assembly cloning kit (New England Biolabs) was used according to the manufacturer’s guidelines.

### Pseudovirus production

Pseudoviruses were produced as previously described by co-transfecting HEK293T cells with expression plasmids for HIV-1 Env or MuLV Env with the *env*-deficient HIV-1 backbone vector pSG3ΔEnv plasmid using FuGENE 6 transfection reagent (Promega).^75^ Cell culture supernatants containing pseudoviruses were harvested after 48 h and filtered through a 0.45 µm filter. FBS was added to a final concentration of 20% and pseudoviruses were stored at - 80°C.

### TZM-bl cell neutralization assays

Neutralization assays were performed as previously described.^17,75^ Briefly, dilutions of antibodies, antibody-containing cell culture supernatants, or heat-inactivated EDTA-plasma or serum (heat-inactivated for 15 min at 56°C) were incubated with viruses on 96-well plates at 37°C for 45 min to 90 min, followed by addition of TZM-bl cells at a final concentration of 10^4^ cells per well in 250 µL medium supplemented with 10-11 µg/mL of DEAE-dextran (Sigma). After 48 h of incubation at 37°C and 5% CO_2_, 150 µL of medium was removed, followed by addition of 100 µL of either Bright-Glo luciferase reagent (Promega) or luciferin/lysis-buffer (17 mM IGEPAL, 0.3 mM ATP, 10 mM MgCl2, 0.5 mM Coenzyme A (all Sigma-Aldrich), and 1 mM D-Luciferin (GoldBio) in Tris-HCL). After 2 min incubation at room temperature, 150 µL of cell lysate was transferred to a black assay plate and read on a luminometer (Berthold). Background relative luminescence units from non-infected control cells were subtracted, and 50% and 80% inhibitory concentrations (for antibodies) or dilutions (for serum and EDTA plasma samples) were calculated as concentrations or dilutions resulting in a 50% or 80% signal reduction, respectively, compared to the signal of untreated virus-infected cells using a four-parameter variable-slope nonlinear regression curve in Prism (GraphPad). All samples were tested at least in technical duplicates.

### Viral outgrowth cultures

For expansion of previously generated bulk CD4 T cell outgrowth viruses obtained through co-cultures with healthy donor CD8-depleted T cells,^17^ MOLT-4 CCR5+ cells were inoculated with viral supernatants and cultured in media without geneticin. Culture supernatants were regularly monitored for p24 production using the Alinity i HIV Ag/Ab Combo Assay (Abbott).

After p24 became highly positive, supernatants were harvested, filtered through a 0.45 µm filter, and stored at -80°C before mice challenge.

### HIV-1 envelope nucleotide sequences and phenotypic analysis

Nucleotide sequences were aligned with Multiple Alignment using Fast Fourier Transform (MAFFT) v7.490 using the 200PAM scoring matrix in Geneious (Dotmatics).^76^ Phylogenetic relationships were inferred using PhyML v3.3 and the general time-reversible (GTR) model with 1,000 bootstraps.^77^ For phylogenetic analysis of IDC#449- and IDC#548-derived *env* in human plasma samples, MOLT4 CCR5+ cell-expanded bulk outgrowth cultures, or infected mice at baseline, trees were rooted to midpoint using iTOL.^78^ For phylogenetic analysis of IDC#449- and IDC#548-derived *env* with compendium clade B sequences, the phylogenetic tree was rooted to HXB2 using iTOL.^78^ 2022 compendium sequences were retrieved from the HIV Los Alamos National Laboratory Sequence Database (https://hiv.lanl.gov). To visualize mutations relative to HXB2 *env*, sequences were aligned with MAFFT v7.490 using Geneious (Dotmatics) and processed using the Los Alamos National Laboratory Sequence Database highlighter tool.^79^

### Amino acid sequence analysis of HIV-1 Env

SGA-derived Env amino acid sequences were aligned using Clustal Omega in Geneious.^80^ For analysis of rebound viruses in HIV-1_YU2_-infected CD34-humanized NXG mice, HIV-1_YU2_ Env was used as a reference sequence, and all disagreements to the reference were highlighted. For analysis of rebound viruses in mice infected with patient virus isolates, mouse-specific baseline majority consensus sequence were generated and compared with mouse-specific post-rebound sequences. Disagreements to baseline sequences within the same mouse were highlighted and colored based on the presence or absence of the amino acid in baseline sequences in other mice.

For the comparison of patient virus isolate-infected mouse baseline sequences, individual consensus sequences were generated based on a nucleotide level MAFFT alignment of SGA-derived Env sequences, using a 75% threshold. Consensus sequences were then translated and aligned to the HXB2 Env sequence using Clustal Omega.

## QUANTIFICATION AND STATISTICAL ANALYSIS

To determine the correlation between 1-18-LS RibobNAb- and 1-18 RibobNAb-containing cell culture supernatant, the Pearson correlation coefficient was calculated in Prism (GraphPad). Spearman’s rank correlation coefficients for the correlation between 1-18-LS RibobNAb serum or plasma concentrations and the serum or plasma HIV-1 neutralizing activity were calculated in Prism (GraphPad). Geometric mean neutralization titers after 1-18-LS RNA-LNP or 1-18-LS recombinant protein injection as well as baseline viral loads between individual mouse treatment groups were compared using a two-tailed Whitney-Mann test (when comparing two groups) or a Kruskall-Wallis test (when comparing three groups) in Prism. Longitudinal comparison of trough antibody levels after repeated injection of 1-18-LS RNA-LNP or 1-18-LS recombinant protein were compared using a mixed-effects model with restricted maximum likelihood (REML) in Prism. Circulating human CD45⁺ cell frequencies in CD34-humanized NRG mice prior to experimental initiation and at study termination were compared using a Wilcoxon signed-rank test in Prism. Viral load changes between groups in HIV-1_YU2_-infected CD34-humanized NRG mice were compared between groups using a Kruskal–Wallis test followed by Dunn’s multiple-comparisons test in Prism. Serum or plasma antibody concentrations below the lower limit of quantification (LLOQ) were imputed as 0.5x LLOQ. For calculations of the log_10_ changes in viremia compared with baseline and depending on the experiment-specific LLOQ, viral loads below 155, 243, or 387 copies/mL were assigned a value of 154, 242, or 386 copies/mL, respectively.

## DATA AND CODE AVAILABILITY

- HIV-1 *env* sequences will be deposited in the GenBank database.
- This paper does not report a custom code.
- Anti-idiotypic antibody against 1-18/1-18-LS, RNA, and LNP-formulated RNA are available from BioNTech to academic, non-commercial researchers with a completed materials transfer agreement upon reasonable request. Requests for these reagents should be directed to Valentin Le Douce.
- Any additional information required to reanalyze the data reported in this work is available from the corresponding authors upon request.

## ACKNOWLEDGMENTS

We thank Chiara Hornung, Anna Schmitt, and the staff of the Animal Care Facility Weyertal at the University of Cologne for facilitating humanized mouse experiments; Lutz Gieselmann for support with single genome amplification; Dominik Aschemeier and Nikolai Grahn for support with HIV-1 pseudovirus cloning; and Acuitas Therapeutics, Inc. for the LNP technology. TZM-bl cells were obtained through the NIH AIDS Reagent Program (ARP) from John C. Kappes and Xiaoyun Wu,^81^ and MOLT-4 CCR5+ cells were obtained through the ARP from M. Baba, H. Miyake, and Y. Iizawa.^82^ P.S. is supported by the German Research Foundation (DFG-Emmy Noether Program, 495793173), and P.S. and H.G. are supported by the Else Kröner-Fresenius-Stiftung. Work in the Humanized Mouse Core Cologne (HMCC) is supported by the German Center for Infection Research (DZIF) (to F.K.). Funding for this study was provided by BioNTech SE.

## AUTHOR CONTRIBUTIONS

Conceptualization, S.S., F.T., R.J., U. Ş., F.K., V.L.D., and H.G.; Methodology, S.S., F.T., U.E., C.R.S., S.K., F.K., V.L.D., and H.G.; Investigation, S.S., R.J., F.T., J.N., A.M., L.F., I.G., N.U., C.M., J.K., R.S., and H.G.; Resources, J.P.B., P.S.; Formal Analysis, S.S., M.S.S., and H.G.; Visualization, S.S., F.T., R.J., A.M., J.P.B., C.M., V.L.D. and H.G.; Writing - Original Draft, S.S. and H.G.; Writing – Review & Editing, F.T., R.J., J.N., C.R.S., S.K., and V.L.D.; Funding Acquisition, F.K.; Supervision, F.T., F.K., V.L.D., and H.G. All authors read and approved the final version of the manuscript.

