## Supplemental Figures for "Durable suppression of viremia by lipid nanoparticle-formulated RNA encoding for a highly potent HIV-1 neutralizing antibody"

### Figure S1

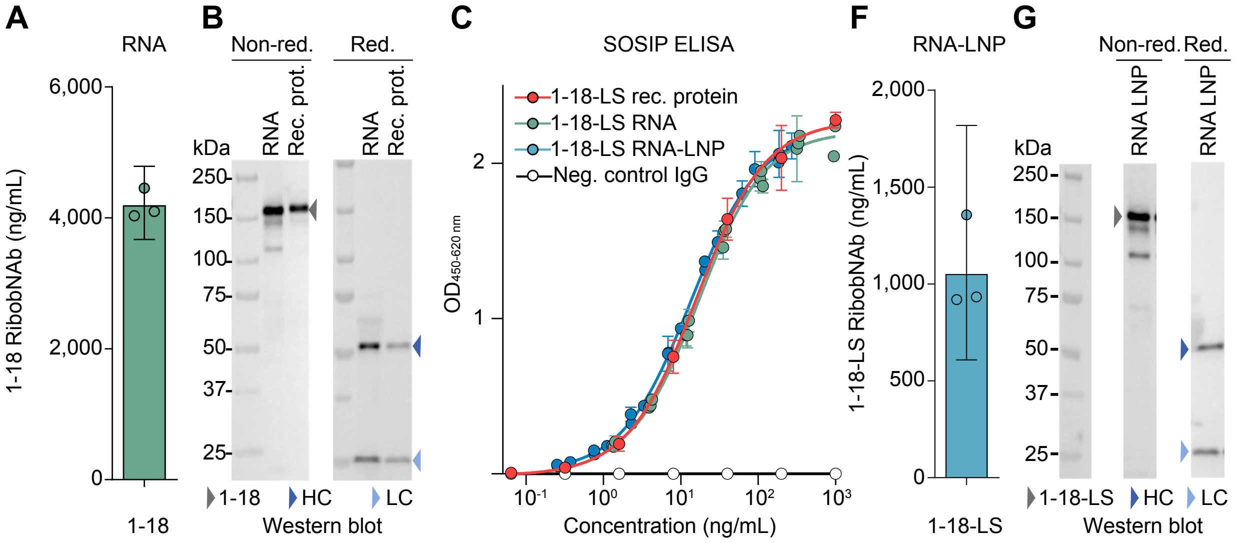


## D

| HIV-1 pseudovirus: HIV-1 strain | Clade | 1-18 protein $IC_{50}$ | 1-18 protein $IC_{80}$ | 1-18-LS RibobNAb $IC_{50}$ | 1-18-LS RibobNAb $IC_{80}$ | $IC_{50}$ FC | $IC_{80}$ FC |
|---|---|---|---|---|---|---|---|
| T255-34 | 02_AG | 0.058 | 0.269 | 0.100 | u.d. | 1.725 | n.a. |
| CNE30 | 08_BC | 0.147 | 0.424 | 0.153 | u.d. | 1.040 | n.a. |
| Ce1176_A3 | C | 0.298 | u.d. | 0.338 | u.d. | 1.132 | n.a. |

<0.1 | 0.1-0.5 | 0.5-1.0 $IC_{50}/IC_{80}$ (µg/mL)

<0.5 | 0.5-2 | >2 Fold change (FC) compared to 1-18 rec. protein

## E

| HIV-1 pseudovirus: HIV-1 strain | Clade | 1-18 rec. protein $IC_{50}$ | 1-18 rec. protein $IC_{80}$ | 1-18-RibobNAb $IC_{50}$ | 1-18-RibobNAb $IC_{80}$ | $IC_{50}$ FC | $IC_{80}$ FC |
|---|---|---|---|---|---|---|---|
| C1080_c3 | 01_AE | 0.136 | 0.479 | 0.151 | 0.267* | 1.110 | 0.556 |
| CNE19 | 07_BC | 0.021 | 0.081 | 0.027 | 0.082 | 1.292 | 1.021 |
| CNE30 | 08_BC | 0.147 | 0.424 | 0.151 | u.d. | 1.024 | n.a. |
| Q23.17 | A1 | 0.036 | 0.107 | 0.034 | 0.098 | 0.947 | 0.915 |
| 6540.v4.c1 | A1C | 0.014 | 0.052 | 0.012 | 0.046 | 0.880 | 0.871 |
| RHPA4259.7 | B | 0.032 | 0.136 | 0.036 | 0.116 | 1.128 | 0.855 |
| TRJO4551.58 | B | 0.127 | 0.445 | 0.115 | 0.256* | 0.906 | 0.576 |
| TRO.11 | B | 0.035 | 0.107 | 0.029 | 0.085 | 0.820 | 0.791 |
| 246F C1G | C | 0.012 | 0.044 | 0.018 | 0.056 | 1.478 | 1.278 |
| Du156.12 | C | 0.038 | 0.134 | 0.084 | 0.214 | 2.243 | 1.599 |
| ZM249M.PL1 | C | 0.041 | 0.177 | 0.054 | 0.160 | 1.325 | 0.902 |
| MuLV (non-HIV control) | - | n.d. | n.d. | n.d. | n.d. | - | - |

<0.1 | 0.1-0.5 | 0.5-1.0 $IC_{50}/IC_{80}$ (µg/mL)

<0.5 | 0.5-2 | >2 Fold change (FC) compared to 1-18 rec. protein

**Supplemental Figure 1. RNA- and RNA-LNP-mediated antibody expression *in vitro*.**
(**A**) 1-18 RibobNAb levels in HEK293T/17 cell culture supernatant after 1-18 heavy and light chain RNA co-transfection. Circles indicate different transfections; bar and error bars show geometric mean and 95% confidence interval, respectively. (**B**) Western blot analysis of 1-18 RibobNAb antibody chain expression in HEK293T/17 cell culture supernatant after 1-18 heavy and light chain RNA transfection compared with 1-18 recombinant protein (rec. prot.). Left and right panels show molecular weight ladder as well as non-reducing (non-red.) and reducing conditions, respectively. Arrows indicate 1-18 antibody, heavy chain (HC), and light chain (LC). (**C**) HIV-1 Env BG505 DS-SOSIP.664 ELISA of 1-18-LS recombinant protein and 1-18-LS RibobNAb in HEK293T/17 cell culture supernatants after RNA transfection or RNA-LNP lipofection. Symbols and error bars indicate arithmetic means and standard deviations, respectively (three transfections measured in technical duplicates for RibobNAbs; recombinant protein tested in four replicates). (**D**) HIV-1 pseudovirus neutralization of 1-18 recombinant protein and 1-18-LS RibobNAb in RNA-transfected HEK293T/17 cell culture supernatants. For supernatants, values represent the mean of triplicates. FC, fold change; u.d., undetermined (due to insufficient antibody concentration in supernatants); n.a., not available. (**E**) HIV-1 pseudovirus neutralization of 1-18 recombinant protein and 1-18 RibobNAb in RNA-transfected HEK293T/17 cell culture supernatant. For supernatants, values represent the mean of triplicates except when indicated by asterisk (in which cases IC80 could only be determined for one or two out of three replicates due to insufficient antibody concentration in supernatants). n.d., not detected at minimum concentrations of 0.30 µg/mL (1-18 protein) or 0.28 µg/mL (1-18 RibobNAb). Other abbreviations as in (D). (**F**) 1-18-LS RibobNAb levels in HEK293T/17 cell culture supernatant after lipofection with 1-18-LS-encoding RNA-LNP. Circles indicate different transfections; bar and error bar show geometric mean and 95% confidence interval, respectively. (**G**) Western blot analysis of 1-18-LS RibobNAb antibody chain expression in HEK293T/17 cell culture supernatant after RNA-LNP lipofection. Left, center, and right panels show molecular weight ladder, non-reducing (non-red), and reducing conditions, respectively. Arrows indicate 1-18-LS antibody, heavy chain (HC), and light chain (LC).

#### Figure S2

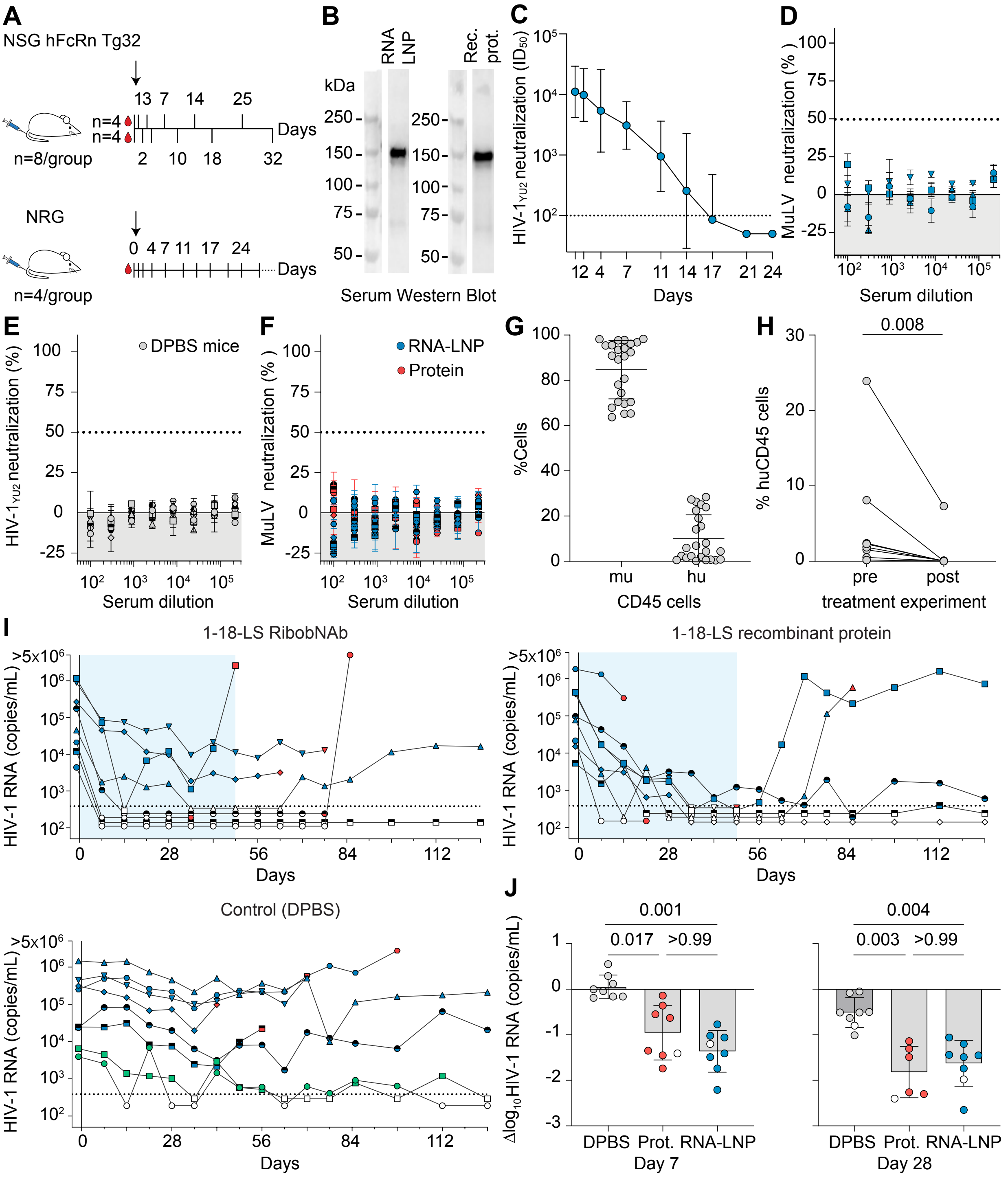


**Supplemental Figure 2. Pharmacokinetics, RibobNAb expression, neutralizing activity, and viral loads in NRG mice.**
(**A**) Experimental scheme for single injection experiments. NSG hFcRn Tg32 mice were analyzed in two parallel groups. (**B**) Representative NSG hFcRn Tg32 mouse serum non-reducing Western blot for human IgG 24 h after 30 µg 1-18-LS RNA-LNP (left) or 100 µg 1-18-LS recombinant protein (rec. prot.) i.v. injection (right). (**C**) Serum HIV-1 YU2 pseudovirus neutralization in NRG mice following a 30 µg 1-18-LS RNA-LNP i.v. injection. Symbols and error bars indicate geometric mean $ID_{50}$s (fifty-percent inhibitory dilutions) and 95% confidence intervals, respectively (*n*=4). Dashed line indicates lowest tested serum dilution. (**D**) Serum murine leukemia virus (MuLV) pseudovirus neutralization in NRG mice one day after a 30 µg 1-18-LS RNA-LNP i.v. injection. Circles and error bars indicate arithmetic mean neutralization and standard deviation of two technical replicates per mouse, respectively (*n*=4). Dashed line indicates 50% neutralization. (**E**) Day 35 serum HIV-1 YU2 pseudovirus neutralization in DPBS-treated HIV-$1_{YU2}$-infected CD34-humanized NRG mice (*n*=8). Symbols indicate different mice; error bars and dashed line as in (D). (**F**) Serum MuLV pseudovirus neutralization in HIV-$1_{YU2}$-infected CD34-humanized NRG mice on day 28, 35, or 42 of weekly 30 µg 1-18-LS RNA-LNP or 500 µg 1-18-LS protein i.v. injections. Symbols indicate different mice; error bars and dashed line as in (D). (**G**) Peripheral blood murine (mu) and human (hu) CD45+ cells 9-13 weeks after humanization (8-12 weeks before HIV-1 challenge) in CD34-humanized NRG mice included in HIV-$1_{YU2}$ infection experiment. Lines and error bars indicate arithmetic means and standard deviation, respectively. (**H**) Human CD45+ cells amongst leukocytes in longitudinally measured NRG mice (8-21 weeks before HIV-1 challenge and 34-47 weeks later) (*n*=8). Groups were compared with a two-tailed Wilcoxon signed-rank test. (**I**) HIV-1 RNA plasma copies in HIV-$1_{YU2}$-infected CD34-humanized NRG mice. Dashed lines indicate HIV-1 RNA lower limit of quantification (LLOQ), white symbols indicate viral loads <LLOQ, and red symbols indicate last measurement in mice that died during the course of the experiment. Shaded areas indicate treatment period (weekly i.v. injections of 500 µg 1-18-LS protein, 30 µg 1-18-LS RNA-LNP, or DPBS). (**J**) $Log_{10}$ viral load changes compared with baseline (day –1) in $HIV_{YU2}$-infected CD34-humanized NRG mice at day 7 (left) and day 28 (right). Bars and error bars indicate arithmetic mean and standard deviation, respectively (*n*=6-8). White symbols indicate viral loads <LLOQ. Viral load changes were compared using a Kruskal–Wallis test followed by Dunn's multiple-comparisons test.

### Figure S3

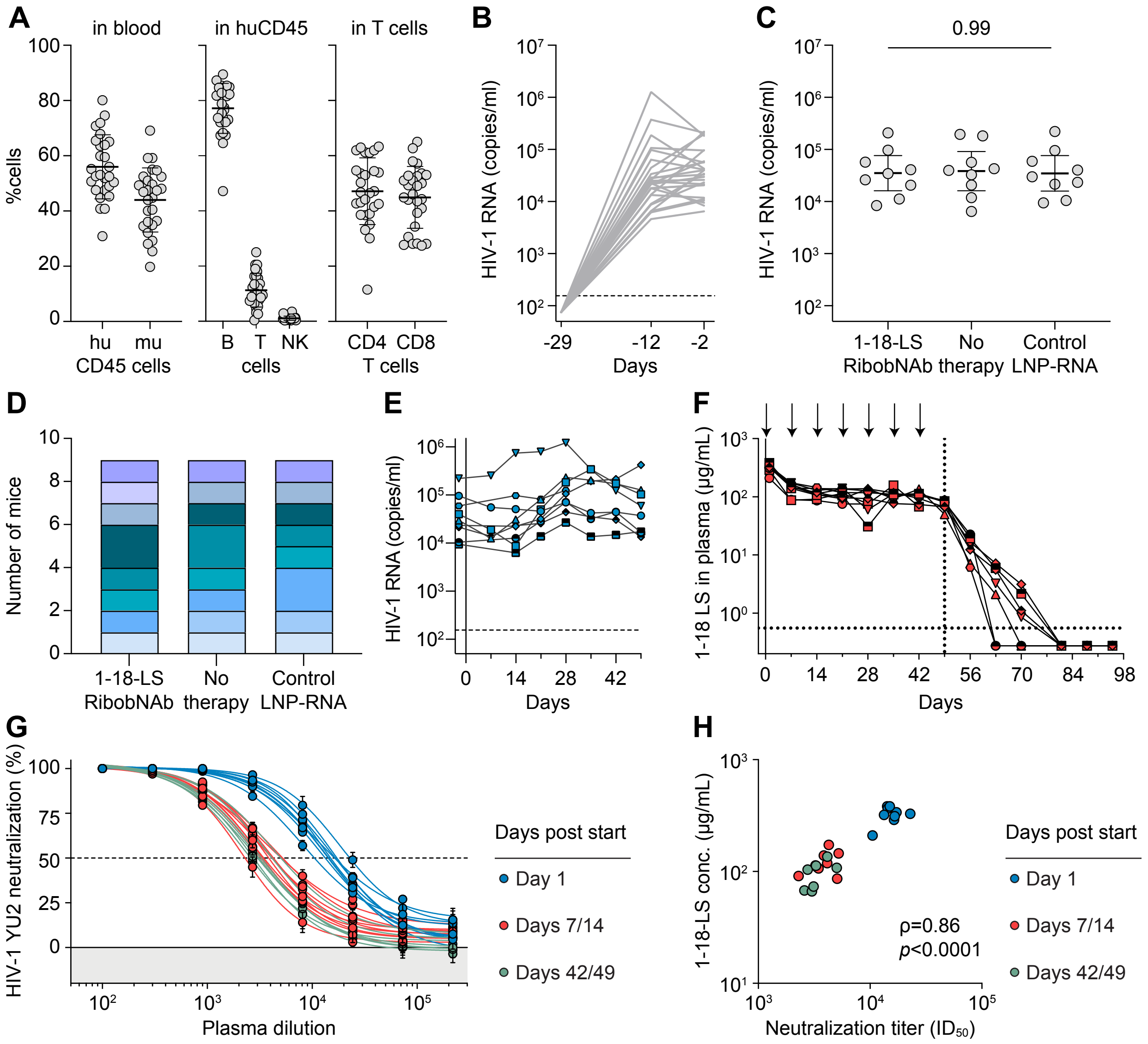


**Supplemental Figure 3. HIV-1$_{YU2}$-infected humanized NXG mice and 1-18-LS RibobNAb levels in vivo.**
(**A**) Human immune cell reconstitution in CD34-humanized NXG mice included in HIV-1$_{YU2}$ infection experiment. Parameters were determined 14-15 weeks after humanization (4-8 weeks prior to viral challenge). Lines and error bars indicate arithmetic means and standard deviation, respectively. (**B**) Plasma viral load trajectories after HIV-1$_{YU2}$ challenge in CD34-humanized NXG mice up to baseline of treatment experiment. Dashed line indicates lower limit of quantification. (**C**) Comparison of baseline HIV-1 plasma loads in 1-18-LS RNA-LNP-treated mice and control groups. Lines and error bars indicate geometric mean and 95% confidence intervals, respectively. Groups were compared using a Kruskal-Wallis test. (**D**) Distribution of CD34 hematopoietic stem cell donors per group of HIV-1$_{YU2}$-infected mice with each color indicating a distinct donor. (**E**) Plasma viral loads in HIV-1$_{YU2}$-infected mice treated weekly with RNA-LNP control constructs. Dashed line indicates the lower limit of quantification (155 copies/mL). (**F**) Plasma 1-18-LS RibobNAb levels in HIV-1$_{YU2}$-infected CD34-humanized NXG mice determined by anti-idiotype ELISA. Arrows indicate intravenous injections of 30 µg 1-18-LS RNA-LNP. Vertical and horizontal dashed lines indicate end of treatment period and lower limit of quantification, respectively. (**G**) Plasma HIV-1 YU2 pseudovirus neutralization in longitudinally followed 1-18-LS RNA-LNP-treated HIV-1$_{YU2}$-infected CD34-humanized NXG mice ($n$=7-9 per time point). Symbols and error bars indicate arithmetic means and standard deviation of two technical replicates per mouse, respectively. (**G**) Correlation of 1-18-LS RibobNAb plasma levels determined by ELISA and plasma HIV-1$_{YU2}$-neutralizing activity in 1-18-LS RNA-LNP-treated mice. $\rho$ indicates Spearman's correlation coefficient determined using all longitudinal data points.

#### Figure S4

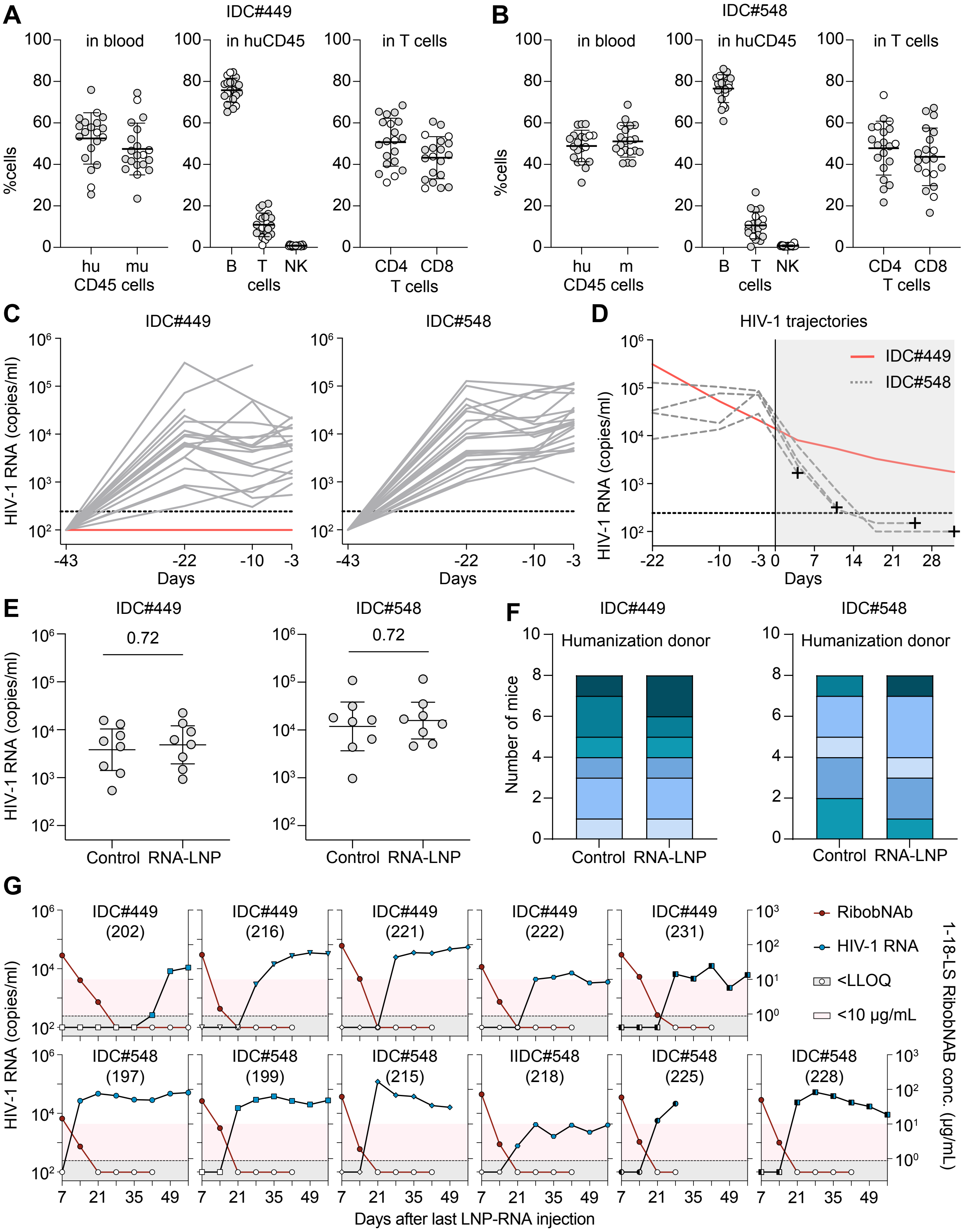


**Supplemental Figure 4. Humanized mice infected with patient virus isolates.**
(**A**) and (**B**) Human immune cell reconstitution parameters in CD34-humanized NXG mice infected with (A) IDC#449- or (B) IDC#548-derived viral isolates. Parameters were determined 9-12 weeks after humanization (13 weeks prior to viral challenge). Lines and error bars indicate arithmetic means and standard deviation, respectively. White circles indicate mice not included in ART interruption experiment (uninfected, *n*=1; incomplete suppression, *n*=1; died before ART initiation *n*=2, died during ART but showed reduction of viremia, *n*=4). (**C**) Plasma viral load trajectories after challenge of CD34-humanized NXG mice with IDC#449- or IDC#548-derived viral isolates up to treatment experiment baseline. Lines indicate individual mice and red line indicates mouse not developing viremia (*n*=1). Dashed lines indicate lower limit of quantification (LLOQ). (**D**) Plasma viral loads in IDC#449-derived isolate-infected CD34-humanized NXG mouse not virologically suppressed during ART (not included in treatment interruption experiments; red line; *n*=1) and IDC#548-derived isolate-infected mice that died during ART with reduced viremia (dashed lines; *n*=4). Area shaded in grey, symbols, and dashed lines indicate ART treatment period, time of death, and LLOQ, respectively. (**E**) Comparison of baseline HIV-1 plasma loads in mice infected with IDC#449- or IDC#548-derived viral isolates and subsequently undergoing ART interruption. Lines and error bars indicate geometric mean and 95% confidence intervals, respectively. Groups were compared using a two-tailed Mann-Whitney test. (**F**) Distribution of CD34 hematopoietic stem cell donors per group of mice infected with IDC#449- or IDC#548-derived viral isolates with colors indicating distinct donors. (**G**) Plasma viral loads and 1-18-LS RibobNAb concentrations after 1-18-LS RNA-LNP treatment interruption. Dashed lines indicate LLOQs for HIV-1 RNA (243 copies/mL) and 1-18-LS RibobNAb concentrations (0.88 µg/mL). White symbols indicate viral loads <LLOQ and areas shaded in red indicate range for antibody levels <10 µg/mL.

#### Figure S5

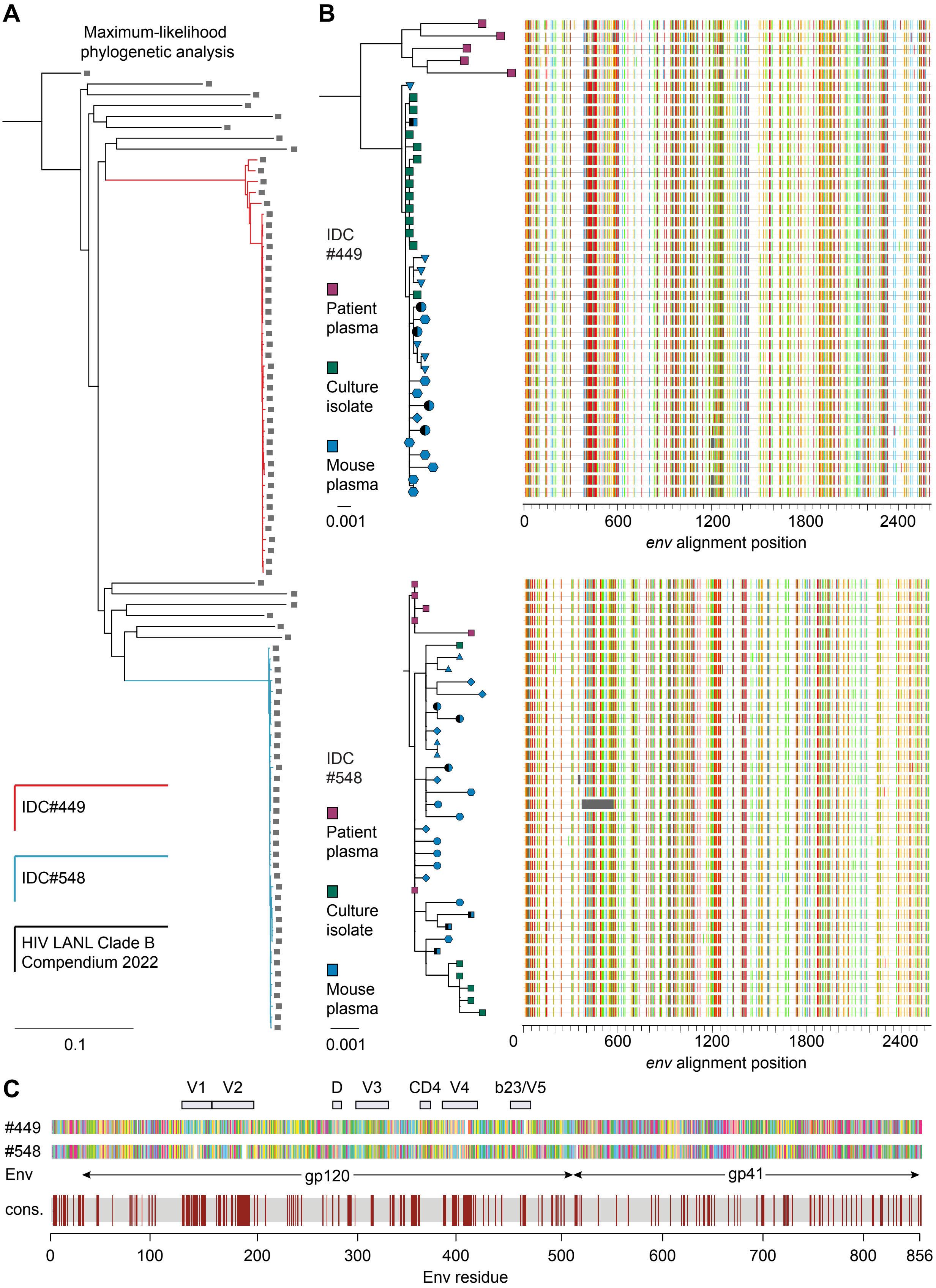


**Supplemental Figure 5. HIV-1 sequence analysis.**
(**A**) Phylogenetic analysis of IDC#449- and IDC#548-derived *env* nucleotide sequences identified by single genome amplification (SGA) in human plasma samples, MOLT-4 CCR5+ cell-expanded bulk outgrowth cultures, or infected mice at baseline, as well as clade B compendium alignment sequences retrieved from the HIV Los Alamos National Laboratory Sequence Database. Bar indicates number of nucleotide substitutions per site. (**B**) Phylogenetic analysis of IDC#449- (top) and IDC#548-derived (bottom) *env* sequences identified by SGA in human plasma samples, MOLT-4 CCR5+ cell expanded bulk outgrowth cultures, or infected mice at baseline with corresponding nucleotide sequences (right). Colored lines indicate nucleic acid mutations relative to the HXB2 reference strain env sequence. Bars indicate number of nucleotide substitutions per site. (**C**) Amino acid sequence alignment of SGA-derived Env consensus sequences obtained at baseline in CD34-humanized NXG mice infected with IDC#449- and IDC#548-derived virus. Bottom graph highlights amino acid sequence differences between the two consensus sequences in red, with residue numbering based on the HXB2 reference strain.
